# A comprehensive and high-quality collection of *E. coli* genomes and their genes

**DOI:** 10.1101/2020.09.21.293175

**Authors:** Gal Horesh, Grace Blackwell, Gerry Tonkin-Hill, Jukka Corander, Eva Heinz, Nicholas R. Thomson

## Abstract

*Escherichia coli* is a highly diverse organism which includes a range of commensal and pathogenic variants found across a range of niches and worldwide. In addition to causing severe intestinal and extraintestinal disease, *E. coli* is considered a priority pathogen due to high levels of observed drug resistance. The diversity in the *E. coli* population is driven by high genome plasticity and a very large gene pool. All these have made *E. coli* one of the most well-studied organisms, as well as a commonly used laboratory strain. Today, there are thousands of sequenced *E. coli* genomes stored in public databases. While data is widely available, accessing the information in order to perform analyses can still be a challenge. Collecting relevant available data requires accessing different sources, where data may be stored in a range of formats, and often requires further manipulation, and processing to apply various analyses and extract useful information. In this study, we collated and intensely curated a collection of over 10,000 *E. coli* and *Shigella* genomes to provide a single, uniform, high-quality dataset. *Shigella* were included as they are considered specialised pathovars of *E. coli*. We provide these data in a number of easily accessible formats which can be used as the foundation for future studies addressing the biological differences between *E. coli* lineages and the distribution and flow of genes in the *E. coli* population at a high resolution. The analysis we present emphasises our lack of understanding of the true diversity of the *E. coli* species, and the biased nature of our current understanding of the genetic diversity of such a key pathogen.

**Author Notes:** All supporting data have been provided within the article or through supplementary data files. All supporting code is provided in the git repository https://github.com/ghoresh11/ecoli_genome_collection.

**Significance as a BioResource to the community:** As of today, there are more than 140,000 *E. coli* genomes available on public databases. While data is widely available, collating the data and extracting meaningful information from it often requires multiple steps, computational resources and expert knowledge. Here, we collate a high quality and comprehensive set of over 10,000 *E. coli* genomes, isolated from human hosts, into a set of manageable files that offer an accessible and usable snapshot of the currently available genome data, linked to a minimal data quality standard. The data provided includes a detailed synopsis of the main lineages present, including their antimicrobial and virulence profiles, their complete gene content, and all the associated metadata for each genome. This includes a database which enables the user to compare newly sequenced isolates against the assembled genomes. Additionally, we provide a searchable index which allows the user to query any DNA sequence against the assemblies of the collection. This collection paves the path for many future studies, including those investigating the differences between *E. coli* lineages, following the evolution of different genes in the *E. coli* pan-genome and exploring the dynamics of horizontal gene transfer in this important organism.

**Data Summary:**

1. The complete aggregated metadata of 10,146 high quality genomes isolated from human hosts (doi.org/10.6084/m9.figshare.12514883, File F1).
2. A PopPUNK database which can be used to query any genome and examine its context relative to this collection (Deposited to doi.org/10.6084/m9.figshare.12650834).
3. A BIGSI index of all the genomes which can be used to easily and quickly query the genomes for any DNA sequence of 61 bp or longer (Deposited to doi.org/10.6084/m9.figshare.12666497).
4. Description and complete profiling the 50 largest lineages which represent the majority of publicly available human-isolated *E. coli* genomes (doi.org/10.6084/m9.figshare.12514883, File F2). Phylogenetic trees of representative genomes of these lineages, presented in this manuscript, are also provided (doi.org/10.6084/m9.figshare.12514883, Files tree_500.nwk and tree_50.nwk).
5. The complete pan-genome of the 50 largest lineages which includes:
  a. A FASTA file containing a single representative sequence of each gene of the gene pool (doi.org/10.6084/m9.figshare.12514883, File F3).
  b. Complete gene presence-absence across all isolates (doi.org/10.6084/m9.figshare.12514883, File F4).
  c. The frequency of each gene within each of the lineages (doi.org/10.6084/m9.figshare.12514883, File F5).
  d. The representative sequences from each lineage for all the genes (doi.org/10.6084/m9.figshare.12514883, File F6).

## Introduction

*E. coli* is a globally distributed, highly diverse organism with a very large gene pool [1–3]. While some variants of *E. coli* are found in the guts of healthy individuals, in animals and in the environment, others cause severe intestinal and extraintestinal life-threatening disease [4]. The diversity between *E. coli* strains is driven by high genome plasticity; genes are regularly gained and lost, leading to high variability in gene content between lineages and isolates [2,5–7]. The combination of these factors: a large gene pool, genome plasticity, global distribution and ubiquity across niches, make *E. coli* an important genetic storehouse for the spread and wider dissemination of genes, including those that confer resistance and virulence. Indeed, *E. coli* has been designated a priority pathogen by the World Health Organisation due to its high levels of drug resistance [8]. Therefore, *E. coli* is a highly relevant organism to study in today’s world, with the increasing spread of antimicrobial resistance, and for understanding the emergence of new, globally disseminated, bacterial pathogens of relevance to human and animal health.

Eight pathogenic variants of *E. coli*, termed “pathotypes”, have been defined based on their site of infection and by distinguishing phenotypic and molecular markers [4]. These are broadly divided into diarrheagenic pathotypes which infect the gastrointestinal tract, and extra-intestinal variants, termed ExPECs, which infect other bodily sites, most notably the urinary tract and the blood. The diarrheagenic pathotypes include Enteropathogenic *E. coli* (EPEC), Enterotoxigenic *E. coli* (ETEC), Enterohaemorrhagic *E. coli* (EHEC), Enteroaggeragive *E. coli* (EAEC), Enteroinvasive *E. coli* (EIEC), Diffusely Adherent *E. coli* (DAEC) and Adherent Invasive *E. coli* (AIEC) [4]. *Shigella* is defined as a separate genus consisting of four different species, *S*.

*sonnei, S. flexneri, S. boydii* and *S. dysenteriae*, for clinical and historical reasons: however, lineages of all *Shigella* species fall within the *E. coli* species phylogeny. Based on molecular definitions, they can be considered diarrheagenic *E. coli [9,10]* with *Shigella* often classified as an EIEC, as they are clinically and diagnostically similar [4]. EPECs, ETECs and *Shigella* are prevalent in the developing world where they cause fatal diarrhea among infants and children [11,12]. ETECs, EAECs and *Shigella* are the most common causes for travellers’ diarrhea [13]. EHECs are the only diarrheagenic *E. coli* that are cause for concern in developed countries as their major reservoir is in the gastrointestinal tracts of cattle [14,15]. EHEC infections cause severe diarrhea, and complications of an infection can cause Haemolytic Uraemic Syndrome (HUS), a life-threatening condition which can lead to kidney failure [4,14].

The transition from non-pathogenic or non-antimicrobial resistant variants of *E. coli* to pathogenic, or antimicrobial resistant, is primarily driven by horizontal gene transfer, through the acquisition of virulence factors or resistance genes on plasmids and other mobile genetic elements [4,16–19]. The availability of thousands of *E. coli* genomes in public databases provides the opportunity to examine the *E. coli* lineages and their gene pool on a scale and resolution that was not previously possible. Here, we collated over 10,000 *E. coli* and *Shigella* isolate genomes, collected from a combination of publications and public databases, and assembled and annotated the entire collection to a high quality. *Shigella* were included as they are phylogenetically part of the *E. coli* species, and are referred to as *E. coli* throughout. We provide all the aggregated associated metadata, a database to query newly sequenced genomes against the assemblies and a searchable index to query a DNA sequence of interest. Additionally, we characterised the most common lineages present in this dataset, including their resistance and virulence profiles. Finally, we defined the complete gene content of these lineages, enabling many future studies examining the biological differences between the lineages and unravelling routes of gene movement in the population.

## Methods

### Data collection

A collection of 18,156 *E. coli* (including *Shigella*) genomes, isolated from human hosts, were downloaded and curated to create a final collection of 10,146 genomes as summarised in Figure 1. For an initial collection of human *E. coli* genomes for which complete metadata is available, whole genome sequences were downloaded from NCBI using genome accessions from publications (detailed in Supplementary File F1 and in the Data Bibliography). The complete metadata was extracted directly from these publications and these were combined. These genomes were supplemented to include other genomes available from public databases, not associated with publications, for which only partial associated metadata was available. These were predominantly sourced from Enterobase and from Public Health England Routine surveillance bioproject (PRJNA315192), downloaded on September 17th, 2018 [3]. As public read repositories also contain pre-publication data, all publicly available genomes were filtered to include only those for which explicit approval was obtained for use by the submitter.

**Figure 1:**
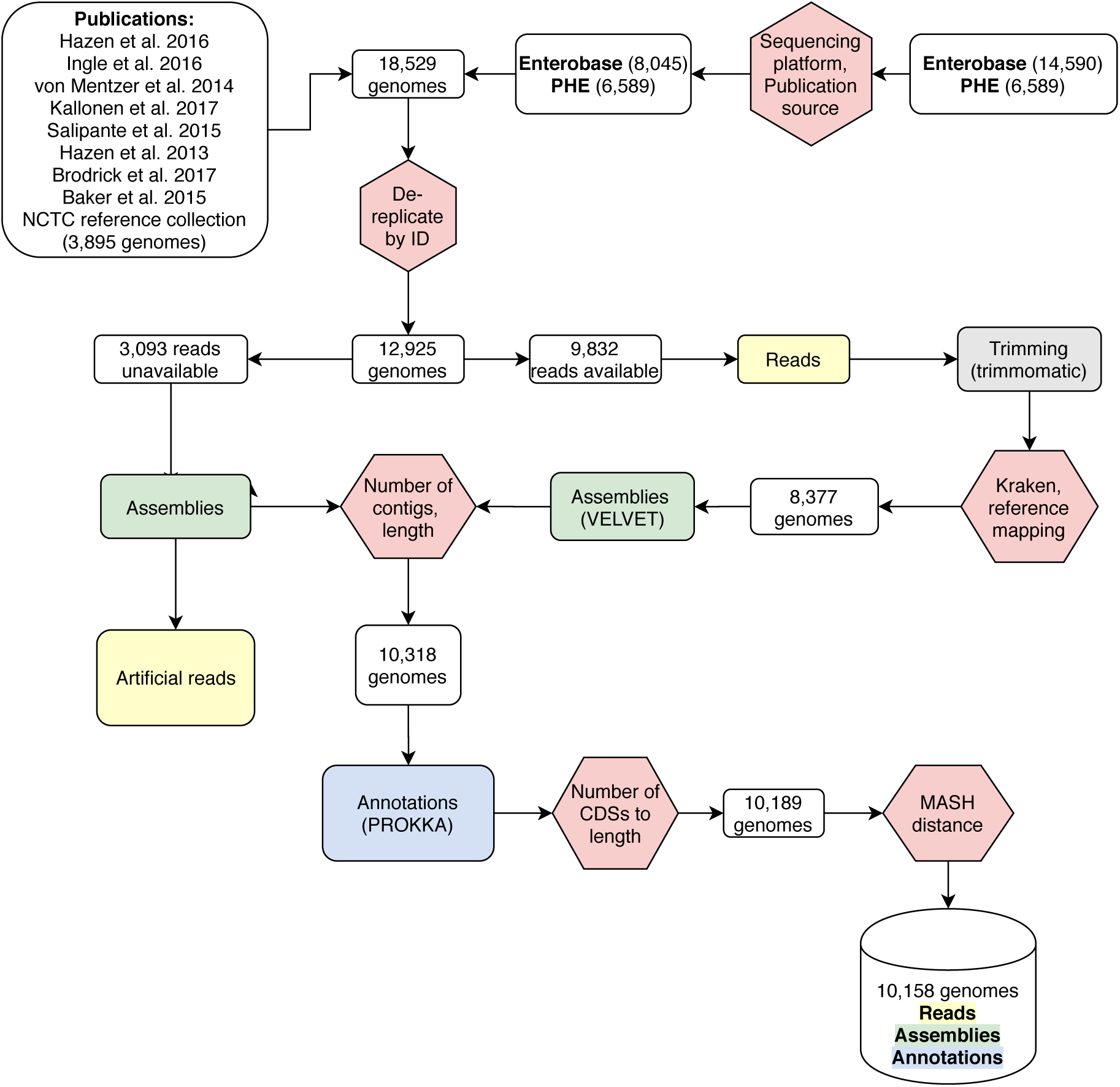
Workflow for constructing the genome collection. Steps taken to obtain a curated, comprehensive and high-quality collection of genomes which include reads, assemblies and annotation files for each included genome. Quality-control steps are in red hexagons, numbers in white squares indicate the number of genomes remaining after each QC step.

#### Reads

Reads were downloaded from the Sequence Read Archive using fastq-dump (v2.9.2). Reads which had been sequenced by Illumina were trimmed using trimmomatic (v0.33) [20] with the *TruSeq3-PE-2* adaptors, a minimum length of 36 bp, and parameters LEADING=10, TRAILING=10, SLIDING WINDOW=4:15 and quality encoding Phred33. When reads were unavailable (3,093 genomes), assemblies were shredded into artificial reads using the script available at https://github.com/sanger-pathogens/Fastaq.

Kraken (v0.10.6) was used on the reads to determine what organism had been sequenced [21]. If fewer than 30% of reads were assigned to *E. coli* or *Shigella* spp., the genome was removed (200 genomes, based on a distribution of these values, Supplementary Figure S1). Reads were also mapped to an *E. coli* reference strain cq9 (GCF_003402955.1) and quality-control (QC) statistics were calculated. Samples were removed (1,255 genomes) according to the distributions of QC values across all reads (Percentage of reads mapped to the reference >60%, percentage of bases mapped that were mismatches was >0.03, percentage of heterozygous SNPs<3%, Supplementary Figure S1).

#### Assembly

Reads were assembled by VELVET (v1.2.09) [22] using the prokaryotic assembly pipeline (v2.0.1) with default setting [23]. Assembled genomes were filtered to remove those with more than 600 contigs or those that had a total combined contig length of less than 4 Mb or larger than 6 Mb (1,152 genomes, based on a distribution of these values, Supplementary Figure S1).

Mash distances were calculated between all the assemblies [24]. Mash uses a minimised database of k-mers, i.e. words of size *k*, to represent each genome (based on the Minhash sketch). Mash returns the proportion of shared k-mers, the Jaccard distance, between every two genomes as a measure of their genomic distance. A network was constructed so that every genome is represented in a node and two genomes were connected only if their Mash distance was smaller than 0.04 (equivalent to 96% Average Nucleotide Identity (ANI)) [24]. Isolates from the same species should have an ANI of approximately 95-96%, i.e. Mash distance smaller than [25]. Therefore, genomes were removed (189 genomes) if they were disconnected from the largest connected component which should represent the *E. coli* and *Shigella* species.

#### Coding sequences

Predicted coding sequences (CDSs) were predicted using Prokka with a custom training file (v1.5, available at doi.org/10.6084/m9.figshare.12514883). Prodigal (v2.6) was trained using a random selected set of 100 genomes from the entire dataset using the “prodigal.py” script available in Panaroo [26,27]. The training file was used as the input for Prokka to predict the CDSs in the entire dataset. All the genomes were then annotated using the same standardised training properties defined in the training file. There was a linear relationship between the size of the genome and the number of genes called. Genomes which deviated from linear correlation by 500 genes were removed (Supplementary Figure S1).

### Constructing the BIGSI index

Each assembly was converted to a non-redundant list of *k*-mers through the construction of De Bruijn graphs (k=31) using mccortex v1.0 [28]. All assemblies had between 105 and 106 unique *k*-mers. The parameters chosen for the BIGSI index were h=1 and m=28000000, as detailed in the berkeleyDB config file (available at doi.org/10.6084/m9.figshare.12666497, file config_10K_00.yaml) and following steps were performed using BIGSI (https://github.com/iqbal-lab-org/BIGSI, checkout:12966cacefc14354ee3c42dc2852917475567a0d) [29]. A single hash function (h=1) was applied to each *k*-mer and each assembly was stored as a fixed length (m=28000000) Bloom filter (bit-vector). To reduce the overall build time of the index, individual Bloom filters were merged in batches of 500 into matrices using ‘bigsi merge_blooms’ command, where the input ‘--from_files’ was a tab separated file where the first column provides the absolute path to the bloom filter and the second is the assembly name. These merged blooms files were then used to build the BIGSI index using ‘bigsi large_build’ command where the provided ‘from_file’ input was a file that contains two columns, separated by tab, where the first column details the absolute path to the merged bloom matrices and the second contains all the corresponding assemblies in that merged bloom file, separated by commas. The BIGSI index of the assemblies in this resource, index10k, can be found at doi.org/10.6084/m9.figshare.12666497.

### Multi-locus sequence typing

The sequence type (ST) for each genome was determined by running “mlst_check” (https://github.com/sanger-pathogens/mlst_check) according to the Achtman MLST scheme downloaded from PubMLST on Jan 22nd, 2019 [30]. *Shigella* are included in the Achtman *E. coli* MLST scheme.

### Defining lineages using PopPUNK

Population Partitioning Using Nucleotide K-mers (PopPUNK) (v. 1.1.3) was used to group the assemblies into PopPUNK clusters or lineages [31]. PopPUNK uses Mash, a k-mer based whole genome comparison approach, to infer the pairwise core and accessory distances between every two assemblies. The database was constructed with parameters k-min=18, k-max=30 and step_size=3, as these values produced the correct line fit for estimating the core and accessory distances, as detailed in https://poppunk.readthedocs.io/en/latest/troubleshooting.html#kmer-length. The estimated core and accessory distances between the assemblies were clustered using a two-dimensional Gaussian mixture model (GMM) to identify cut-offs for the within lineage core and accessory distances. The model fitting was applied using six different values of total number of clusters for the GMM (K= 5, 8, 11, 14, 17 and 20). The scores generated by PopPUNK for all these values were compared. A value of K=11 was chosen as it had the overall lowest entropy, i.e. highest confidence in assigning each distance to a cluster, and comparably high overall score. PopPUNK then constructs a network between all assemblies where each node is an assembly and two assemblies are connected only if their core and accessory distance is below the “within lineage core” and “within lineage accessory” distances. All assemblies which are connected to each other in this network are defined as a lineage.

### Phylogenetic analysis

The core gene phylogeny was inferred from the core gene alignment generated using Roary for each lineage [32], and a tree from the SNPs in the core gene alignment, extracted using SNP-sites [33] (v2.3.2), was constructed using FastTree [34]. Treemer (v0.3)[35] was used to select ten genomes from each lineage as representatives of that lineage (Supplementary Table S1). Similarly, Treemer was used to choose a single representative genome from each of the 50 lineages to generate a tree containing only 50 genomes. In both cases, the core gene phylogeny was inferred from the SNPs of the core gene alignment generated using Roary on the representative genomes [32]. A maximum likelihood tree from the informative SNPs, chosen using SNP-sites [33] (v2.3.2), was constructed using RAxML (v8.2.8)[36] with 100 bootstrap replicates.

### Phylogroup assignment

ClermonTyping (v1.4.1) was used to assign the *E. coli* phylogroup of the 500 representative *E. coli* genomes [37]. ClermonTyping uses an *in-silico* PCR approach of marker genes, following the Clermont phylotyping scheme presented in [38]. This is supplemented by a Mash-based mapping to a curated collection of *E. coli* genomes, for which the phylogroup is known. A lineage was assigned to the phylogroup according to the most common phylogroup assignment of the ten representative strains. The exception was Lineage 10 which was assigned to phylogroup D by ClermonTyping as the marker gene *arpA* was not detected in the *in-silico* PCR using primer ArpAgpE, however, the assignment did not correspond with the phylogeny and this was corrected to phylogroup E.

### Identification of antimicrobial and virulence genes

A collection of antimicrobial resistance (AMR) genes were obtained from ResFinder (https://bitbucket.org/genomicepidemiology/resfinder_db/src/master/, downloaded on 06.03.19) [39]. Virulence genes were downloaded from the Virulence Finder Database (https://bitbucket.org/genomicepidemiology/virulencefinder_db/src, downloaded 24/08/18). Read files of genomes (real where available or otherwise, artificially generated from the assemblies) were queried for the presence of these known AMR or virulence genes using ARIBA (v2.14) with default settings [40]. A gene was marked as present only if 80% of the entry sequence in the database was covered, otherwise it was marked as absent.

### Pathotype assignments

Pathotypes were assigned according to the presence of specific marker virulence genes according to the pathotype-associated markers presented in Table 1 in [41], refined by the source of isolation: if the source of isolation was blood or urine the assignment was ExPEC; if any variant of shiga-toxin was present the assignment was STEC; if *eae* was present the assignment was aEPEC/EPEC; if both shiga-toxin and *eae* were present the assignment was EHEC; if either *aatA, aggR* or *aaiC* were present the assignment was EAEC; if *est* or *elt* were present the assignment was ETEC; if *ipaH9*.*8* or *ipaD*, characteristic of the invasive virulence plasmid pINV, were present the assignment was EIEC. A pathotype was assigned to a lineage if at least half of the isolates of the lineage were assigned to the same pathotype. *Shigella* lineages were assigned *Shigella* as their pathotype.

### Pan-genome analysis

A pan-genome analysis using Roary [32] was applied on each lineage separately using the default identity cutoff of 0.95, with paralog splitting disabled [32]. The outputs of the pan-genome analysis of each lineage were combined to generate a final collection of gene clusters of the entire dataset in the following steps:

1. Gene cluster definitions, from the Roary analysis within each lineage, were assumed to be the best approximation of the representation of the genes that are well-defined within a closely related group of genomes. Note that each gene cluster has multiple members (sequences) from that lineage (Supplementary Figure S2, Step 1). A representative sequence was chosen for each gene cluster as the sequence that had the modal length within that gene cluster. If there was no mode, a sequence with the median length was chosen.
2. A pan-genome analysis using Roary was applied on all lineages in an all-against-all manner using an identity threshold of 0.95 and with paralog splitting disabled, leading to a total of 1,081 Roary analyses. This generated gene clusters for each possible lineage pair. Note that, similar to Step 1, each gene cluster can have multiple members (sequences), but this time from both lineages used in each respective comparison (Supplementary Figure S2, Step 2).
3. A “combined Roary graph” was constructed, with the gene clusters from the original Roary outputs from Step 1 (a Roary analysis on a single lineage only) as nodes (Supplementary Figure S2, Step 3).
4. Gene cluster of lineage A was connected to a gene cluster of lineage B if there was a gene clustering in their combined Roary analysis (step 2) where i) 80% of the members of the gene cluster of A were in the new combined clustering, and ii) 80% of the members of the gene cluster of B were also in the combined clustering (Supplementary Figure S2, Step 4).
5. The following corrections were applied to add or remove connections between gene clusters in the combined Roary graph (Supplementary Figure S2, Step 5):
  a. Density based clustering groups data points based on their density in space, while assuming that data points which belong to the same group are in a region of a high density and are separated from another group by a region of low density.
  b. The distance metric used for density based clustering was the proportion of shared edges (Jaccard index) between every two nodes in the combined Roary graph. This identified spurious connections between genes which were not supported by most pairwise Roary analyses (Supplementary Figure S2). This was applied using the “dbscan” method of the python package sci-kit learn [42] with parameters epsilon=0.5 and min_samples=6. Connections between a gene cluster of Lineage A and a gene cluster of Lineage B that did not belong to the same dbscan cluster were removed.
  c. To correct for under-splitting, all representative sequences of each gene cluster of the combined Roary graph were aligned to each other using mafft (v7.310) [43] with default settings. If the alignment of two sequences showed more than 20% mismatches along the length of the longer sequence, the connection between them in the combined Roary graph was removed (See Supplementary Figure S2, Step 5b).
  d. To correct for over-splitting, the representative sequences of all the gene clusters of the original Roary outputs were aligned to each other using blastp (version 2.9). Representative sequences which were more than 95% identical over 80% of their length were merged (See Supplementary Figure S2, Step 5c).
6. Following corrections, the connected components of the combined Roary graph were the final set of gene clusters in the entire dataset (Supplementary Figure S2, Step 6).

File F6 in doi.org/10.6084/m9.figshare.12514883 contains the representative sequences from the original Roary outputs (Step 1) for each gene in the final gene clusters (Step 6).

### Statistical analysis

Statistical analyses were performed in R (v3.3+). Ape (v5.3) [44] and ggtree (v1.16.6) [45] were used for phylogenetic analysis and visualisation. The ggplot2 (v3.2.1) package was used for plotting [46]. All scripts used in the analysis are available at: https://github.com/ghoresh11/ecoli_genome_collection

## Results

18,156 *E. coli* genomes, isolated from human hosts, were collected from a variety of sources and required multiple steps which are detailed in the Methods section and summarised in Figure 1. *Shigella*, which are phylogenetically part of the *E. coli* species, were also included and are referred to as *E. coli* throughout. In short, genome identifiers from publications where complete metadata was available were collected, and combined with identifiers of genomic data from public databases for which only limited metadata was available. Genomes were downloaded, assembled and their CDSs were predicted and annotated. Importantly, to ensure the accuracy of the data, multiple QC measures were applied, reducing the initial dataset and thereby ensuring a final collection of high-quality genomes (Figure 1B). Only genomes for which we received explicit approval to be used by the submitter were kept, removing any doubts regarding the ability to use this data for high resolution analyses. The curated high-quality final genome collection comprises 10,146 genomes on which all the subsequent analysis was performed. This makes this dataset unique as it can be used as a reliable, well-described and curated reference for the diversity of the majority of publicly available human-isolated *E. coli* genomes.

The vast majority of available *E. coli* genomes are from developed countries, collected in surveillance in clinical settings. The clinical samples are mostly generated by agencies which conduct regular investigations of *E. coli* isolates in outbreaks and routine surveillance programmes. These include Public Health England (PHE) (5,207 genomes), the Food and Drug Administration (FDA) (883 genomes) and the Centers for Disease Control and Prevention (CDC) (561 genomes) (Supplementary Figure S3). This explains the bias in the available genomic data with 70% and 15% of the original samples originating from the United Kingdom (UK) or the United States (US), respectively. The remaining genomes originated mostly from other countries in Europe, with only a small fraction of genomes being currently available from Asia, Africa, South America or Oceania.

38% of the samples considered here were taken from faeces, blood and urine. The remaining samples were recorded unknown or other “human sources” (File F1). Isolates from Africa and Asia were exclusively from faecal samples, whereas isolates from Europe and North America included those causing both intestinal and extraintestinal disease (Supplementary Figure S3). Where available, the pathotype description was as described in the original publication. Within these isolates, the representation of diarrheal disease-causing *E. coli* pathotypes, EPECs and ETECs, was very low with only 3% and 2% of the genomes belonging to these pathotypes, respectively.

### Six STs represent more than 50% of the genomes in the collection

Multi-locus sequence typing is based on the variation of seven housekeeping genes, the combination of which define a sequence type or ST. A total of 993 different known STs were identified in the collection. 87 STs (9%) alone accounted for 80% of the isolates (Supplementary Figure S4). Six STs, 11, 131, 73, 10, 95 and 21, accounted for 50% of the isolates included here. 790 STs (∼80% of the STs) were represented by five isolates or fewer. Many of the former represent important STs linked to human disease. For instance, ST11 (30% of all genomes) is associated with EHEC serotype O157:H7, a major foodborne pathogen that can be contracted by eating contaminated foods, specifically beef products, as it lives in the colon of cattle and is an important cause of HUS in humans [14]. The collection also includes STs of non-O157 EHECs, including STs 17 (2%) and 21 (2%). STs 131 (8%), 73 (4%), and 95 (3%) are all STs known to be associated with extra-intestinal disease [47–49]. ST10 (3%) is a broad host range ST, isolates of which have been observed in multiple host species, and include all known *E. coli* pathotypes [50].

### The dataset can be divided into lineages of closely related isolates

As *E. coli* is a highly diverse organism, relying on MLST for subtyping can lead to new ST definitions within a group of closely related isolates due to variation in one of these genes, or otherwise, to connections between unrelated isolates due to recombination. We therefore grouped the genomes into lineages of closely related isolates using a whole-genome based approach. PopPUNK extracts and compares words of size k, named “k-mers”, from whole genomes to measure the deviation in core-gene sequence termed as the “core distance”, and the deviation in gene content, termed as the “accessory distance”, between two genomes [31]. In *E. coli*, the core distance, as estimated by PopPUNK, correlates with the pairwise SNP distance between all the core genes of the two genomes being compared, and the accessory distance correlates with the proportion of shared accessory genes between every two genomes (the Jaccard distance) [31]. Genomes which had both low core and accessory distances were considered to be in the same PopPUNK cluster, defined here as a “lineage”, as they were highly similar in both their core and accessory genomes.

Based on the rules described above this grouping produced 1,154 lineages. As expected the distribution of lineage sizes was similar to that defined by MLST with a few large lineages representing most of the population (Supplementary Figure S4). A single lineage, Lineage 1, contained 34% of all genomes (File F2). This lineage was mostly comprised of ST11, i.e. O157:H7 EHEC. Similarly, Lineage 2 contained 8% of all genomes and consisted mostly of ST131, a global multidrug resistant (MDR) ExPEC lineage. The third largest lineage, Lineage 3, contained 5% of all genomes and mostly consisted of isolates belonging to ST73 (File F2).

### 50 PopPUNK lineages represent more than 75% of the genomes, and are representative of the currently known *E. coli* population structure

We focused the further analysis of the dataset on the 50 lineages which had at least twenty isolates. Together these lineages included 7,693 genomes (76% of the collection) and 271 different STs (27% of those described by this collection).

To examine the population structure and diversity of the 50 lineages largest lineages, the phylogeny was constructed by selecting ten genomes from each lineage that captured most of the diversity of that lineage (See Methods, Supplementary Table S1), leading to a total of 500 genomes representing the dataset. Their core genome was extracted and the phylogenetic tree from the core gene alignment was inferred. The phylogenetic analysis confirmed that PopPUNK separated the genomes into clearly distinct lineages based on their core genome (Figure 2). The exception to this was Lineage 12 which was split into two closely related groups. One group was more closely related to Lineage 28 whereas the other to Lineage 35. The core and accessory distances estimated by PopPUNK showed that indeed, the core distance between PopPUNK Clusters 12, 28 and 35 was low, however, they sufficiently deviated in their accessory gene content to be defined as three distinct PopPUNK lineages.

**Figure 2:**
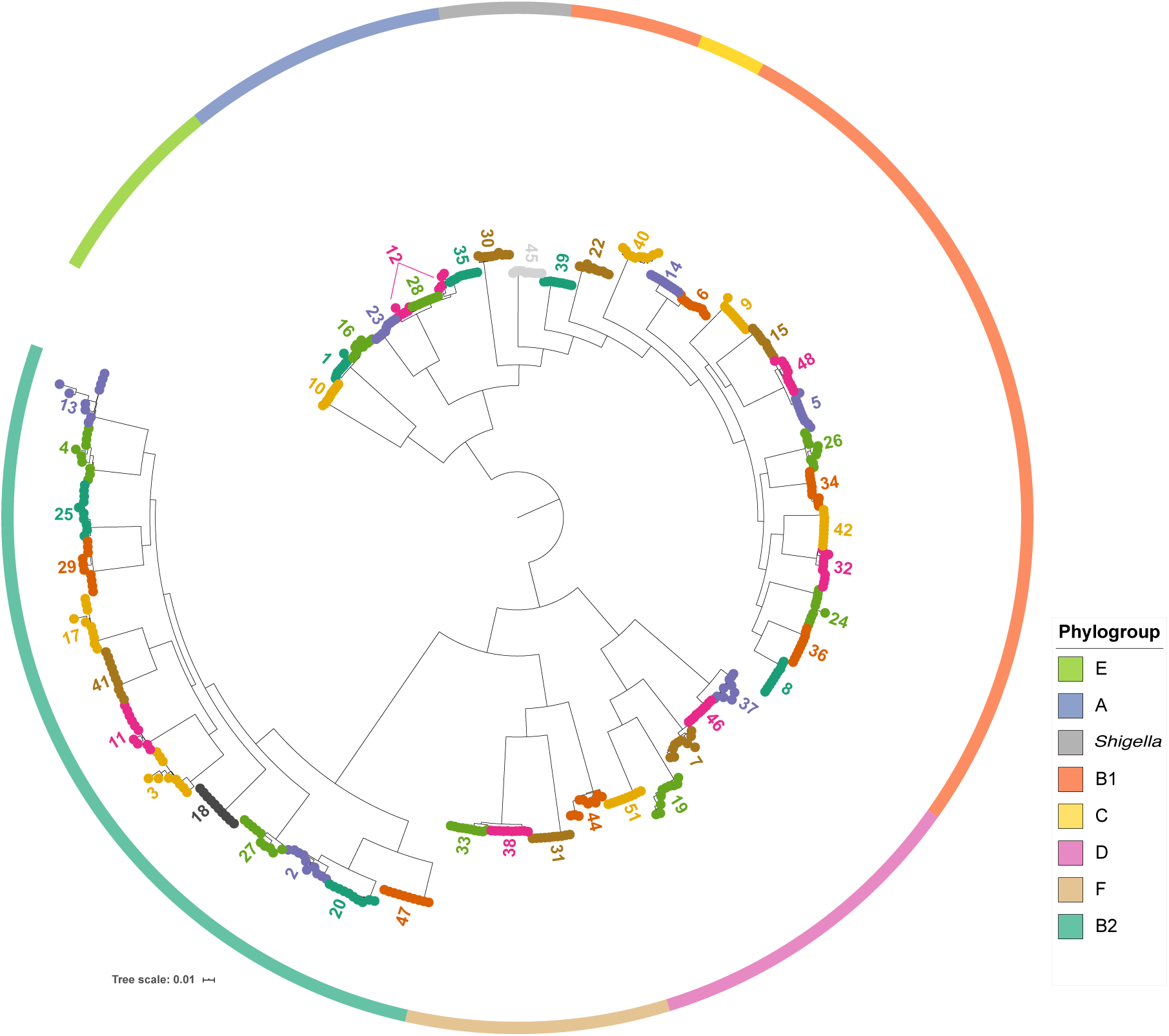
Population structure of the lineages. Core gene phylogeny of 10 representatives from each of the 50 largest PopPUNK lineages, selected using Treemer [35]. Solid coloured outer ring indicates the phylogroup assignment of the representatives of that lineage. The tree was plotted using ITOL [51]. Colours on tips used to distinguish between the PopPUNK lineages.

Population genetics studies on *E. coli* have defined the existence of eight deep-branching phylogenetic groups, termed “phylogroups” (A, B1, B2, D, E, F, C and G) [52–55]. While the collection assembled here is biased towards particular STs and we only included lineages with 20 genomes or more, it is evident from Figure 2 that the collection of genomes spans all *E. coli* phylogroups (18 from B1, 13 from B2, 4 from A, 5 from D, 4 from F, 3 from E, 1 from C and 2 of *Shigella* representing *S. sonnei* (45) and *S. flexneri* (30) and therefore is representative of the known species diversity [38,56].

### Associated metadata shows a consistent source of isolation per lineage

The lineages broadly divide into those enriched for isolates collected from faecal samples and those collected from blood and urine samples (See File F2, Supplementary Figure S5). Only lineages 26, 34 and 48 of the intestinal isolate lineages were enriched for samples collected from Africa and Asia. These lineages mostly represented EPEC and ETEC isolates which had been collected from faecal samples in developing countries as part of the The Global Enteric Multicenter Study (GEMS) collection, in contrast to the other lineages containing faecal samples which include STECs or EHECs were collected in the high income settings [12]. Lineage 12, which consisted of 78% isolates from ST10, was the only lineage that spanned all continents and consisted of all sample types (faecal, blood, urine or unknown).

Where sampling date was available, 39% of the genomes in the collection were collected in the last 10 years. A number of lineages included older, historically important isolates from the Murray collection [57] (Supplementary Figure S5). Notably, Lineage 30, which contains *S. flexneri* isolates, had a higher proportion of isolates collected before 1980 relative to the rest of the collection (Wilcox summed rank test, p<0.05, Bonferroni corrected).

### Lineages vary substantially in their genome size

The number of genes in a single isolate and the size of the genome varied significantly between the lineages (Supplementary Figure S6). The weighted-mean number of genes across all lineages was 4,869 genes and the weighted-mean genome length was 5.2 Mbp. Isolates from the *Shigella* lineages 30 and 45 had the smallest genomes with a genome size of only 4.3 and 4.7 Mbp. Lineages 12, 40 and 48 had the second smallest genome lengths with a mean genome length of ∼4.85Mbp. On the other hand, Lineages 5, 6, 8, 15, and 48, all from phylogroup B1, had a mean of over 5,100 genes per isolate (200 genes more than the dataset mean). The number of predicted genes and genome size were affected by the phylogroup. Lineages in phylogroups E, F and B1 tended to have larger genomes with a few exceptions. Lineages from phylogroup C, B2 and A tended to have smaller genomes. Phylogroup D had a wider range of observed genome sizes.

### Multidrug resistance was predicted for more than half of the isolates in 16 of 50 lineages

A total of 153 known resistance gene alleles were identified in the collection. The number of known resistance genes within each isolate ranged from none to a maximum of 18 in a single isolate, predicted to confer resistance to up to ten different antimicrobial classes (Figure 3A, File F1).

**Figure 3:**
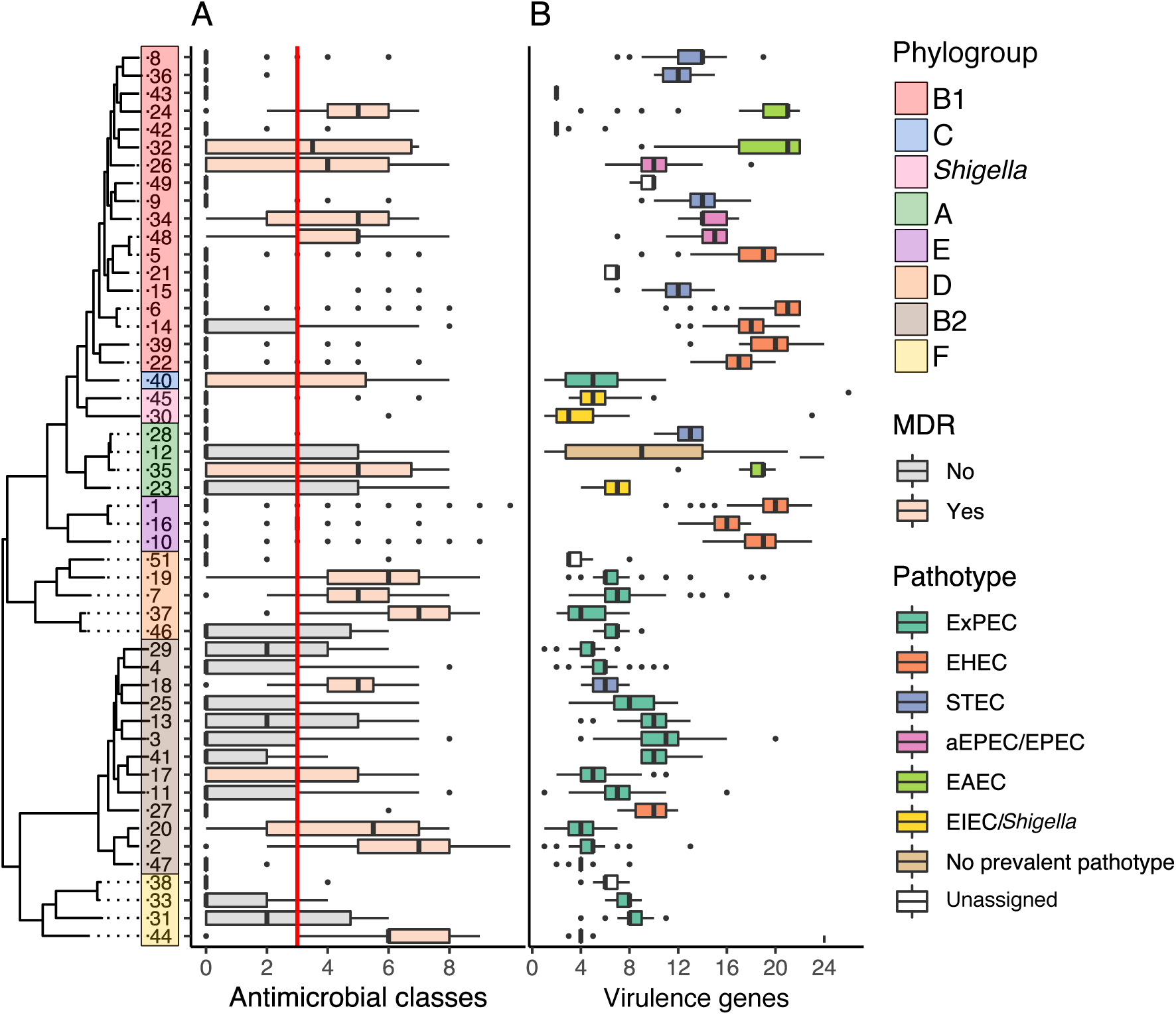
Antimicrobial resistance and virulence profiles of the lineages. **A** Number of predicted antimicrobial classes each isolate is resistant to, based on genetic profile by lineage. Red line indicates threshold for multidrug resistance (predicted resistance to three classes of antimicrobials or more). **B** Number of virulence genes per isolate, by lineage and coloured by the most prevalent predicted pathotype in the lineage. ND = “Not Determined”.

Multidrug resistance in an isolate has been defined as resistance to three classes of antibiotics or more [58]. All but five lineages (Lineages 21, 36, 43, 47 and 49) had at least one isolate which was MDR. We defined an MDR lineage as a lineage where half of the isolates or more were MDR. 16 of the 50 lineages investigated were MDR (Figure 3A, File F2). Importantly, this metric is affected by the sampling bias; lineages are MDR because isolates with clinical significance are being sequenced, and it does not inform on the true diversity of antimicrobial resistance genotypes within these lineages in the *E. coli* population. Indeed, *E. coli* isolated from humans have been shown to possess more resistance genes [59]. Half of these lineages were isolated predominantly from blood and urine samples, i.e. ExPECs (Lineages 2, 20, 44, 40, 17, 7, 37 and 9). These included lineages 2 and 20 which contain isolates of the global ExPEC lineage ST131. Three of the ExPEC MDR lineages belonged to phylogroup D (Lineages 19, 7 and 37). Three other MDR lineages predominantly contained EPEC isolates from the GEMS collection (Lineages 26, 34 and 48) [12,60]. The source of isolation of the remaining five lineages (Lineages 32, 35, 18, 16 and 24) was predominantly unknown (Supplementary Figure S5).

### 43 of 50 lineages are dominated by a single *E. coli* pathotype

Consistent with the collection of *E. coli* isolates being from human hosts and mostly from clinical samples, 439 known virulence genes were observed in our dataset. The isolates had a median of nine known virulence genes in a single genome, with a maximum value of 26 virulence genes present in a single isolate.

A combination of the source of isolation as well as the detection of a set of marker virulence genes were used to find the most prevalent predicted pathotype within each lineage (See Methods). 44 of 50 lineages were identified as predominantly containing one of the *E. coli* pathotypes, i.e. at least half of the isolates of the lineages were predicted to belong to one of the pathotypes (Figure 3B). Lineage 12, which mostly consists of *E. coli* isolates typing as ST10, was the only lineage which contained isolates assigned to multiple different pathotypes with no single dominant pathotype (11% ExPEC, 29% EAEC, 24% EPEC, 9% STEC, 2% EHEC, 1% ETEC, and 24% unassigned). The remaining six lineages which were not assigned an *E. coli* pathotype, predominantly from B1 (21,42,43,49;B1, 38;F, and 51;D), had relatively few virulence genes as well as few AMR genes.

Of the isolates included here Phylogroups B2, F, and D predominantly contained ExPEC isolates. Lineages 27 and 18 were the only lineages in phylogroup B2 which contained 67% EHEC isolates and 33% aEPEC/EPECs (Lineage 27) and 100% STEC isolates (Lineage 18) (Figure 3B). All phylogroup E lineages contained predominantly EHEC isolates. Phylogroups A and B1 had more diversity of pathotypes, containing lineages which were assigned to the range of diarrheagenic pathotypes (EPEC, EHEC, EAEC and EIEC). Lineage 24 of phylogroup B1 contained 38% isolates which were *stx* and *eae* positive. These are isolates of *E. coli* serotype O104:H4 taken from the 2011 German outbreak, which were classified as the convergence of an EHEC and an EAEC [61]. Lineage 40 was the only ExPEC lineage within the B1-C-A clade.

### The final pan-genome includes a total of 55,039 genes

In order to define the gene content of this reference collection, an initial pan-genome analysis was applied to the lineages separately (see Methods), revealing a low gene diversity within Lineages 21, 43 and 49 (Supplementary Figure S7). Therefore, these were not included in the detailed description of the pan-genome of the lineages as the low diversity was linked to these being collected at the same time by the FDA. The outputs of the 47 pan-genome analyses of the remaining lineages were combined in order to provide a description of the gene pool in the entire dataset (see Methods). Briefly, a pairwise pan-genome analysis was applied on all CDSs of every two lineages. The grouping of CDSs in every pairwise pan-genome analysis was examined to determine whether two CDSs from two lineages should be labelled as the same gene in the complete dataset.

A total of 55,039 predicted CDSs were identified in this dataset (Files F3-F6). As there are 47 lineages, and a varying number of isolates per lineage, each gene has a frequency within each of the 47 Lineages (provided in File F5). For instance, the *intA* gene, encoding a prophage integrase, was observed in 20 of the lineages (Figure 4A). In two lineages (6 and 9), it was present in over 95% of isolates, in another 8 lineages it was present in intermediate frequencies (between 15% and 95%) and in the final 10 lineages it was present in fewer than 15% of isolates. In contrast, the gene *wzyE*, a gene involved in antigen biosynthesis, is a core gene which was observed across all lineages in a frequency of over 95% (Figure 4B). Principal component analysis on all the gene frequencies across the lineages showed that the first and second principal components explained 17.93% and 7.49% of the variance and separated the lineages by phylogroup (Figure 4C).

**Figure 4:**
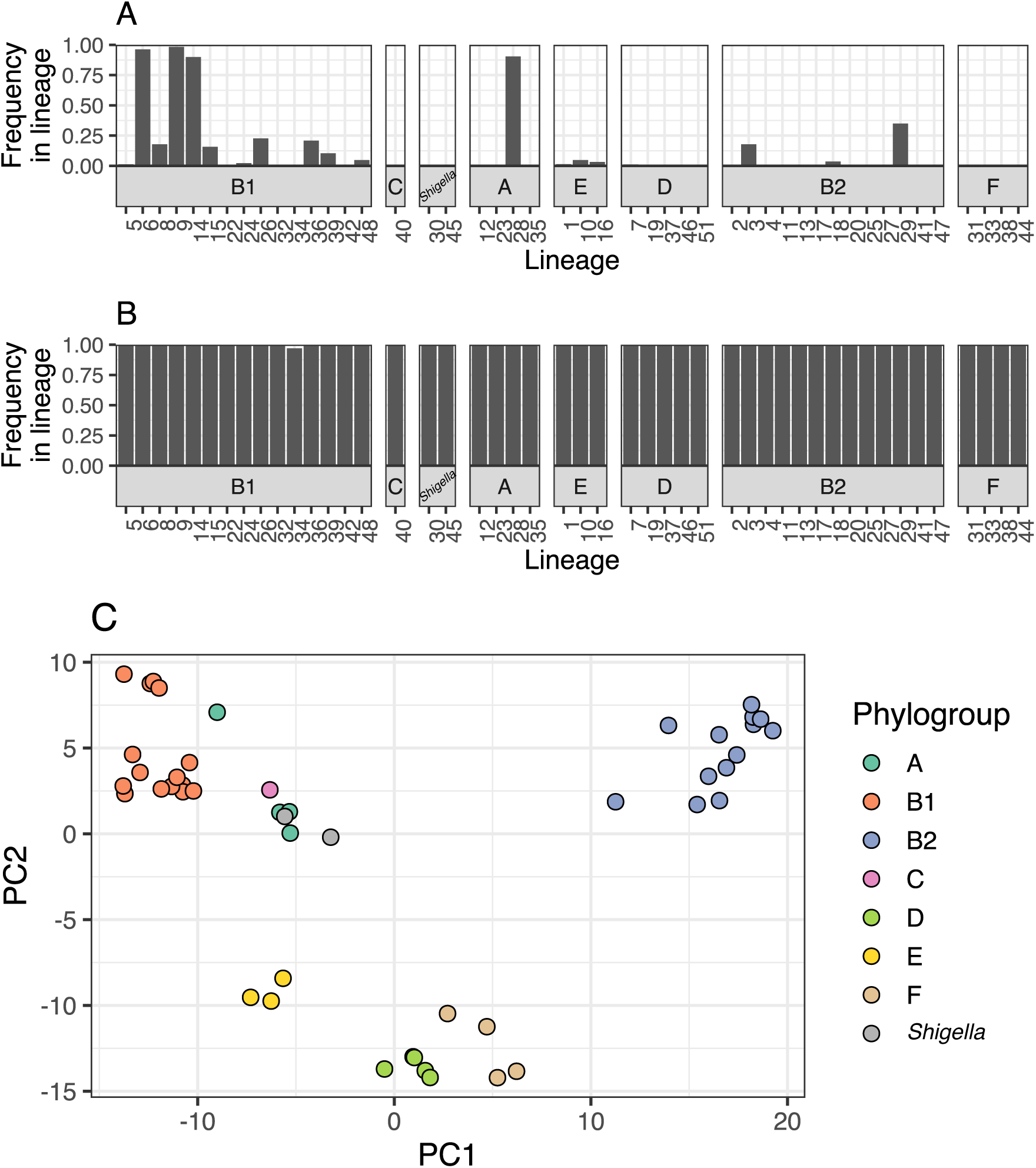
Gene frequencies across the lineages. **A, B:** Examples of the frequencies of two genes across the 47 lineages, stratified by phylogroup. *intA* (**A**) is present in some lineages and is observed in different frequencies across these. *wzyE* (**B**) is a core gene observed in a high frequency across all lineages. **C** PCA plot of the gene frequencies across all lineages, coloured by phylogroup.

## Example usage

### Searching for any DNA sequence in the collection using BIGSI

BIGSI uses a k-mer based approach to query any DNA sequence of 61 bp or greater against all the assemblies of the collection [29]. This can be achieved as follows, using the files provided at doi.org/10.6084/m9.figshare.12666497:

~~~
bigsi search -c config_10K_00.yaml -t 0.8
ATGAAAAACACAATACATATCAACTTCGCTATTTTTTTAATAATTGCAAATATTATCTACA
~~~

Where config_10K_00.yaml provides the config file to the BIGSI index of the assemblies, and 0.8 is the threshold in k-mer similarity (equivalent to 2 mismatches per 100 bps) used to defined a match, and “ATGAAAAACACAATACATATCAACTTCGCTATTTTTTTAATAATTGCAAATATTAT CTACA” is the sequence being used to search the dataset, compiled here using BIGSI. BIGSI will return all the genome identifiers in the collection that have this sequence in at least 80% k-mer similarity. The properties of these genomes can be investigated in File F1. The user will need to ensure the path to the index is correct (“filename:”) in the config_10K_00.yaml file. Please refer to the BIGSI documentation (https://github.com/iqbal-lab-org/BIGSI) for full details.

### Examine the membership of newly sequenced genomes to the lineages in this collection

Newly sequenced genomes can be compared to the lineages in this collection by using the PopPUNK Database provided at doi.org/10.6084/m9.figshare.12650834, as follows:

~~~
poppunk --assign-query --ref-db ecoli_poppunk_db --q-files list_of_genomes.txt --output out
~~~

Where ecoli_poppunk_db is the PopPUNK database provided above, and list_of_genomes.txt is a file containing the list of the new user provided assemblies being queried. A new directory named ‘out’ is automatically created. The file out_clusters.csv will capture the assignment of each assembly to the lineages defined in the PopPUNK database. The properties of these lineages can be examined in files provided in this manuscript F1 and F2. Please refer to the PopPUNK documentation (https://poppunk.readthedocs.io/en/latest/) for full details.

### Examining the distribution of a gene across the species phylogeny

A gene of interest can be identified in the pan-genome presented by using alignment tools like Blast+ [62] or DIAMOND [63] against the pan genome reference file provided (File F3). The distribution of the gene named “intA_1”, a prophage integrase, in this genome collection can be plotted across the phylogeny of the 47 lineages using the frequencies from the provided File F5 (Figure 5A). The phylogeny of the specific sequences of each lineage can be drawn using the sequences provided in File F6 (Figure 5B).

**Figure 5:**
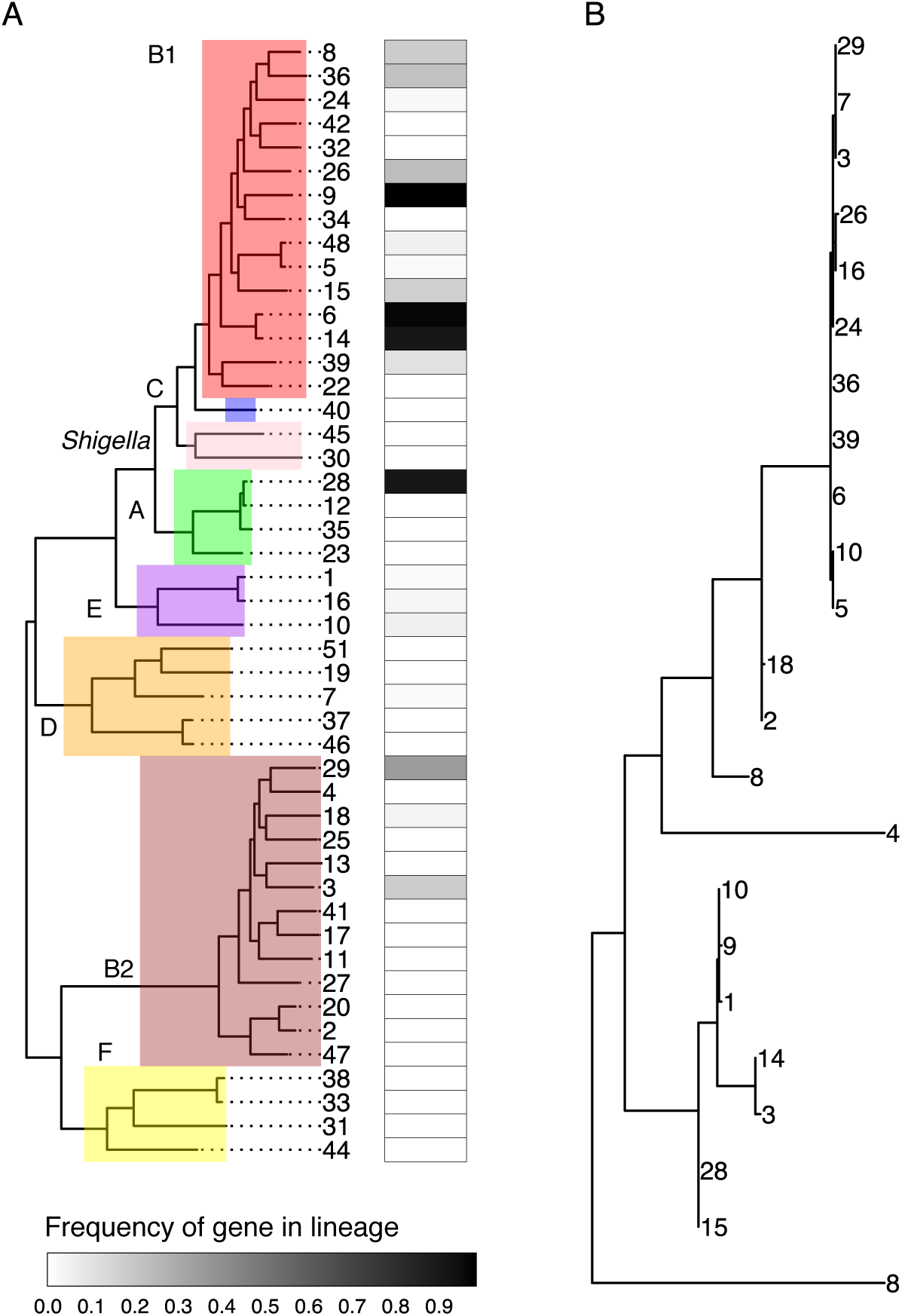
Example usage of the pan-genome to examine the distribution of a single, prophage integrase gene (intA_1). **A** The distribution of a gene can be examined across the species tree using the gene frequencies from File F5. Heatmap indicates the fraction of isolates of a lineage that possess the gene. **B** Phylogenetic tree of the gene sequences from each lineage. Sequences of the gene from panel A in each lineage can be extracted from File F6 to examine the species wide evolution of the gene. Numbers on branch tips indicate the lineage.

## Discussion

We have created a high-quality, extensively curated dataset of over 10,000 *E. coli* and *Shigella* genomes, linked this to resources which enable this dataset to be queried as a single dataset, and have provided several usage examples. Additionally, we have described in detail the properties of the main lineages present in the collection and their gene (predicted CDS) content. We hope that the data provided in this manuscript will make future studies on *E. coli* more accessible to a wider audience and will facilitate the investigation of some of the pressing questions in *E. coli* genetics and evolution.

Aggregating data from diverse sources along with their associated metadata is not trivial but, given the increasing number of data sources and data types, essential. Genome identifiers and data formats across publications and databases do not always match leading to many conversions which are error prone and require knowledge of programming. In addition, computational resources are required in order to apply thousands of assembly and annotation calculations. These are all limiting factors to research. This emphasises the need to build new resources which maintain high quality genome collections where users would more easily be able to both retrieve and apply analyses on large collections. Without such resources, information is widely available, but it is practically only usable for a small proportion of scientists with large resources and computational expertise. Enterobase is a valuable resource which overcomes data accessibility issues by integrating, assembling and analysing the genomic data of specific enteric pathogens from the Sequence Read Archive, while providing researchers with relevant metadata and software [3]. However, as metadata is often associated with a publication, and is not directly linked to the database from which the genome was downloaded, this information is often missing. Even more, describing the gene content by comparing whole genomic datasets is a much harder problem, which cannot realistically be provided in a high quality in an automated manner across increasing dataset sizes. Therefore, studies on *E. coli* in recent years have either been detailed and focused only on a single pathotype [60,64–66] or, when utilising a very large number of genomes, the analyses were limited in their resolution due to the complexity of extracting the information from such large collections [3,67]. Taken together, the collection presented here represents a detailed, high-quality and accessible dataset which will enable researchers to apply comprehensive comparisons in future investigations on *E. coli*. This includes the PopPUNK and BIGSI databases which can be used to query newly sequenced isolates or DNA sequences of interest and examine their diversity relative to this collection.

The analysis presented in this manuscript emphasises our lack of knowledge on the true diversity of this important species, and that we should redirect our efforts towards sampling to understand the diversity which has yet to be studied. The collection we obtained is biased towards *E. coli* lineages which have clinical significance. The vast majority of genomes were available from Europe and North America, such that the pathotypes comprising the dataset are those which predominantly affect these areas. Of 1,154 lineages, there were only 50 which contained at least 20 isolates which were used for defining the gene-content. Sampling should be increased in a directed manner in under-represented areas of the world as well as sampling of non-clinical isolates. Using the PopPUNK database provided in this study, future studies can incorporate new genomes to the dataset provided here and compare *E. coli* isolates from other geographical locations, animals or the environment to the genomes presented here. The PopPUNK database could be expanded and updated in future versions which include these more targeted samples which expand on the diversity presented here.

Biological differences between the lineages were already revealed from the initial descriptions of the lineages presented in this study. There were clear differences in the genome size between the phylogroups and lineages. Higher variability in genome size within a phylogroup or lineage could be an indication of higher rates of gene gain and loss within that lineage. A larger genome size may also help to equip a lineage to survive in a range of niches. These results indicate the importance of this dataset in addressing some important questions regarding the differences between different *E. coli* lineages and gene flow in the *E. coli* population.

## Funding Information

Wellcome Sanger Institute [206194]; Wellcome Sanger Institute Ph.D. Studentship (to G.H.); Wellcome Trust PhD Scholarship Grant [204016 to G.T.H]; ERC grant 742158 to (J.C).

### Acknowledgements

We would like to thank Leopold Parts, Simon Harris, Andres Floto and members of the Thomson Team for useful discussions. We would also like to thank Cinzia Fino for collating genomic identifiers of ExPEC isolates from publications.

## Abbreviations

HGT: Horizontal Gene Transfer
EPEC: Enteropathogenic *E. coli*
ETEC: Enterotoxigenic *E. coli*
EHEC: Enterohaemorrhagic *E. coli*
EAEC: Enteroaggeragive *E. coli*
EIEC: Enteroinvasive *E. coli*;
DAEC: diffusely adherent *E. coli*
AIEC: adherent invasive *E. coli*
ExPEC: extraintestinal *E. coli*
CDS: coding sequence
ST: sequence type
AMR: antimicrobial resistance
PHE: Public Health England
FDA: Food and Drug Administration
CDC: Centers for Disease Control and Prevention
GEMS: Global Enteric Multicenter Study
MDR: multidrug resistant
SNP: Single Nucleotide Polymorphism

## Data bibliography

1. Zhou Z, Alikhan N-F, Mohamed K, Fan Y, Agama Study Group, Achtman M. The EnteroBase user’s guide, with case studies on *Salmonella* transmissions, *Yersinia* pestis phylogeny, and *Escherichia* core genomic diversity. Genome Res. 2020;30: 138–152.

2. Public Health England Routine surveillance Bioproject (PRJNA315192), downloaded on September 17th, 2018

3. Kallonen T, Brodrick HJ, Harris SR, Corander J, Brown NM, Martin V, et al. Systematic longitudinal survey of invasive *Escherichia coli* in England demonstrates a stable population structure only transiently disturbed by the emergence of ST131. Genome Res. 2017. doi: 10.1101/gr.216606.116.

4. Brodrick HJ, Raven KE, Kallonen T, Jamrozy D, Blane B, Brown NM, et al. Longitudinal genomic surveillance of multidrug-resistant *Escherichia coli* carriage in a long-term care facility in the United Kingdom. Genome Med. 2017;9: 70.

5. Salipante SJ, Roach DJ, Kitzman JO, Snyder MW, Stackhouse B, Butler-Wu SM, et al. Large-scale genomic sequencing of extraintestinal pathogenic *Escherichia coli* strains. Genome Res. 2015;25: 119–128.

6. Baker KS, Burnett E, McGregor H, Deheer-Graham A, Boinett C, Langridge GC, et al. The Murray collection of pre-antibiotic era Enterobacteriacae: a unique research resource. Genome Med. 2015;7: 97.

7. von Mentzer A, Connor TR, Wieler LH, Semmler T, Iguchi A, Thomson NR, et al. Identification of enterotoxigenic *Escherichia coli* (ETEC) clades with long-term global distribution. Nat Genet. 2014;46: 1321–1326.

8. Ingle DJ, Tauschek M, Edwards DJ, Hocking DM, Pickard DJ, Azzopardi KI, et al. Evolution of atypical enteropathogenic *E. coli* by repeated acquisition of LEE pathogenicity island variants. Nat Microbiol. 2016;1: 15010.

9. Hazen TH, Sahl JW, Fraser CM, Donnenberg MS, Scheutz F, Rasko DA. Refining the pathovar paradigm via phylogenomics of the attaching and effacing *Escherichia coli*. Proc Natl Acad Sci U S A. 2013;110: 12810–12815.

10. Hazen TH, Donnenberg MS, Panchalingam S, Antonio M, Hossain A, Mandomando I, et al. Genomic diversity of EPEC associated with clinical presentations of differing severity. Nat Microbiol. 2016;1: 15014.

11. Public Health England NCTC 3000 reference collection (https://www.phe-culturecollections.org.uk/collections/nctc-3000-project)

12. Goh KGK, Phan M-D, Forde BM, Chong TM, Yin W-F, Chan K-G, et al. Genome-Wide Discovery of Genes Required for Capsule Production by Uropathogenic *Escherichia coli*. MBio. 2017;8. doi: 10.1128/mBio.01558-17

13. Chen SL, Wu M, Henderson JP, Hooton TM, Hibbing ME, Hultgren SJ, et al. Genomic diversity and fitness of *E. coli* strains recovered from the intestinal and urinary tracts of women with recurrent urinary tract infection. Sci Transl Med. 2013;5: 184ra60.

## References

1. Rasko DA, Rosovitz MJ, Myers GSA, Mongodin EF, Fricke WF, Gajer P, et al. The pangenome structure of *Escherichia coli*: comparative genomic analysis of *E. coli* commensal and pathogenic isolates. J Bacteriol. 2008;190: 6881–6893.

2. Touchon M, Hoede C, Tenaillon O, Barbe V, Baeriswyl S, Bidet P, et al. Organised genome dynamics in the *Escherichia coli* species results in highly diverse adaptive paths. PLoS Genet. 2009;5: e1000344.

3. Zhou Z, Alikhan N-F, Mohamed K, Fan Y, Agama Study Group, Achtman M. The EnteroBase user’s guide, with case studies on *Salmonella* transmissions, *Yersinia pestis* phylogeny, and *Escherichia* core genomic diversity. Genome Res. 2020;30: 138–152.

4. Croxen MA, Law RJ, Scholz R, Keeney KM, Wlodarska M, Finlay BB. Recent advances in understanding enteric pathogenic *Escherichia coli*. Clin Microbiol Rev. 2013;26: 822–880.

5. Welch RA, Burland V, Plunkett G 3rd, Redford P, Roesch P, Rasko D, et al. Extensive mosaic structure revealed by the complete genome sequence of uropathogenic *Escherichia coli*. Proc Natl Acad Sci U S A. 2002;99: 17020–17024.

6. Wirth T, Falush D, Lan R, Colles F, Mensa P, Wieler LH, et al. Sex and virulence in *Escherichia coli*: an evolutionary perspective. Mol Microbiol. 2006;60: 1136–1151.

7. Didelot X, Méric G, Falush D, Darling AE. Impact of homologous and non-homologous recombination in the genomic evolution of *Escherichia coli*. BMC Genomics. 2012;13: 256.

8. Organization WH, Others. Global priority list of antibiotic-resistant bacteria to guide research, discovery, and development of new antibiotics. Geneva: World Health Organization. 2017.

9. Pettengill EA, Pettengill JB, Binet R. Phylogenetic Analyses of *Shigella* and Enteroinvasive *Escherichia coli* for the Identification of Molecular Epidemiological Markers: Whole-Genome Comparative Analysis Does Not Support Distinct Genera Designation. Front Microbiol. 2015;6: 1573.

10. Chattaway MA, Schaefer U, Tewolde R, Dallman TJ, Jenkins C. Identification of *Escherichia coli* and *Shigella* Species from Whole-Genome Sequences. J Clin Microbiol. 2017;55: 616–623.

11. Ochoa TJ, Contreras CA. Enteropathogenic *Escherichia coli* infection in children. Current Opinion in Infectious Diseases. 2011. pp. 478–483. doi: 10.1097/qco.0b013e32834a8b8b

12. Kotloff KL, Nataro JP, Blackwelder WC, Nasrin D, Farag TH, Panchalingam S, et al. Burden and aetiology of diarrhoeal disease in infants and young children in developing countries (the Global Enteric Multicenter Study, GEMS): a prospective, case-control study. Lancet. 2013;382: 209–222.

13. de la Cabada Bauche J, Dupont HL. New Developments in Traveler’s Diarrhea. Gastroenterol Hepatol. 2011;7: 88–95.

14. Nguyen Y, Sperandio V. Enterohemorrhagic *E. coli* (EHEC) pathogenesis. Front Cell Infect Microbiol. 2012;2: 90.

15. Dean-Nystrom EA, Bosworth BT, Moon HW. Pathogenesis of *Escherichia coli* O157:H7 in Weaned Calves. In: Paul PS, Francis DH, editors. Mechanisms in the Pathogenesis of Enteric Diseases 2. Boston, MA: Springer US; 1999. pp. 173–177.

16. Carattoli A. Plasmids and the spread of resistance. Int J Med Microbiol. 2013;303: 298–304.

17. Acheson DW, Reidl J, Zhang X, Keusch GT, Mekalanos JJ, Waldor MK. *In vivo* transduction with shiga toxin 1-encoding phage. Infect Immun. 1998;66: 4496–4498.

18. Dudley EG, Thomson NR, Parkhill J, Morin NP, Nataro JP. Proteomic and microarray characterization of the AggR regulon identifies a pheU pathogenicity island in enteroaggregative *Escherichia coli*. Mol Microbiol. 2006;61: 1267–1282.

19. Pilla G, Tang CM. Going around in circles: virulence plasmids in enteric pathogens. Nat Rev Microbiol. 2018;16: 484–495.

20. Bolger A, Giorgi F. Trimmomatic: a flexible read trimming tool for illumina NGS data. URL http://www.usadellab/org/cms/indexphp. 2014.

21. Wood DE, Salzberg SL. Kraken: ultrafast metagenomic sequence classification using exact alignments. Genome Biol. 2014;15: R46.

22. Zerbino DR, Birney E. Velvet: algorithms for de novo short read assembly using de Bruijn graphs. Genome Res. 2008;18: 821–829.

23. Page AJ, De Silva N, Hunt M, Quail MA, Parkhill J, Harris SR, et al. Robust high-throughput prokaryote de novo assembly and improvement pipeline for Illumina data. Microb Genom. 2016;2: e000083.

24. Ondov BD, Treangen TJ, Melsted P, Mallonee AB, Bergman NH, Koren S, et al. Mash: fast genome and metagenome distance estimation using MinHash. Genome Biol. 2016;17: 132.

25. Kim M, Oh H-S, Park S-C, Chun J. Towards a taxonomic coherence between average nucleotide identity and 16S rRNA gene sequence similarity for species demarcation of prokaryotes. Int J Syst Evol Microbiol. 2014;64: 346–351.

26. Tonkin-Hill, G., MacAlasdair, N., Ruis, C. et al. Producing polished prokaryotic pangenomes with the Panaroo pipeline. Genome Biol 21, 180 (2020). https://doi.org/10.1186/s13059-020-02090-4

27. Hyatt D, Chen G-L, Locascio PF, Land ML, Larimer FW, Hauser LJ. Prodigal: prokaryotic gene recognition and translation initiation site identification. BMC Bioinformatics. 2010;11: 119.

28. Turner I, Garimella KV, Iqbal Z, McVean G. Integrating long-range connectivity information into de Bruijn graphs. Bioinformatics. 2018;34: 2556–2565.

29. Bradley P, den Bakker HC, Rocha EPC, McVean G, Iqbal Z. Ultrafast search of all deposited bacterial and viral genomic data. Nat Biotechnol. 2019;37: 152–159.

30. Page AJ, Taylor B, Keane JA. Multilocus sequence typing by blast from de novo assemblies against PubMLST. J Open Source Softw. 2016;1: 118.

31. Lees JA, Harris SR, Tonkin-Hill G, Gladstone RA, Lo SW, Weiser JN, et al. Fast and flexible bacterial genomic epidemiology with PopPUNK. doi: 10.1101/360917

32. Page AJ, Cummins CA, Hunt M, Wong VK, Reuter S, Holden MTG, et al. Roary: rapid large-scale prokaryote pan genome analysis. Bioinformatics. 2015;31: 3691–3693.

33. Page AJ, Taylor B, Delaney AJ, Soares J, Seemann T, Keane JA, et al. SNP-sites: rapid efficient extraction of SNPs from multi-FASTA alignments. Microb Genom. 2016;2: e000056.

34. Price MN, Dehal PS, Arkin AP. FastTree 2--approximately maximum-likelihood trees for large alignments. PLoS One. 2010;5: e9490.

35. Menardo F, Loiseau C, Brites D, Coscolla M, Gygli SM, Rutaihwa LK, et al. Treemmer: a tool to reduce large phylogenetic datasets with minimal loss of diversity. BMC Bioinformatics. 2018;19: 164.

36. Stamatakis A. RAxML version 8: a tool for phylogenetic analysis and post–analysis of large phylogenies. Bioinformatics. 2014;30: 1312–1313.

37. Beghain J, Bridier-Nahmias A, Le Nagard H, Denamur E, Clermont O. ClermonTyping: an easy-to-use and accurate in silico method for *Escherichia* genus strain phylotyping. Microb Genom. 2018;4. doi: 10.1099/mgen.0.000192

38. Clermont O, Christenson JK, Denamur E, Gordon DM. The Clermont *Escherichia coli* phylo-typing method revisited: improvement of specificity and detection of new phylogroups. Environ Microbiol Rep. 2013;5: 58–65.

39. Zankari E, Hasman H, Cosentino S, Vestergaard M, Rasmussen S, Lund O, et al. Identification of acquired antimicrobial resistance genes. J Antimicrob Chemother. 2012;67: 2640–2644.

40. Hunt M, Mather AE, Sánchez-Busó L, Page AJ, Parkhill J, Keane JA, et al. ARIBA: rapid antimicrobial resistance genotyping directly from sequencing reads. Microb Genom. 2017;3: e000131.

41. Robins-Browne RM, Holt KE, Ingle DJ, Hocking DM, Yang J, Tauschek M. Are *Escherichia coli* Pathotypes Still Relevant in the Era of Whole-Genome Sequencing? Front Cell Infect Microbiol. 2016;6: 141.

42. Pedregosa F, Alexandre Gramfort N, Michel V, Thirion B, Grisel O, Blondel M, et al. Scikit-learn: Machine Learning in Python. J Mach Learn Res. 2011;12: 2825–2830.

43. Katoh K, Standley DM. MAFFT multiple sequence alignment software version 7: improvements in performance and usability. Mol Biol Evol. 2013;30: 772–780.

44. Paradis E, Claude J, Strimmer K. APE: Analyses of Phylogenetics and Evolution in R language. Bioinformatics. 2004;20: 289–290.

45. Yu G, Smith DK, Zhu H, Guan Y, Lam TT. ggtree: an r package for visualization and annotation of phylogenetic trees with their covariates and other associated data. McInerny G, editor. Methods Ecol Evol. 2017;8: 28–36.

46. Wickham H. ggplot2: Elegant Graphics for Data Analysis. Springer; 2016.

47. Kallonen T, Brodrick HJ, Harris SR, Corander J, Brown NM, Martin V, et al. Systematic longitudinal survey of invasive *Escherichia coli* in England demonstrates a stable population structure only transiently disturbed by the emergence of ST131. Genome Res. 2017

48. Brodrick HJ, Raven KE, Kallonen T, Jamrozy D, Blane B, Brown NM, et al. Longitudinal genomic surveillance of multidrug-resistant *Escherichia coli* carriage in a long-term care facility in the United Kingdom. Genome Med. 2017;9: 70.

49. Day MJ, Doumith M, Abernethy J, Hope R, Reynolds R, Wain J, et al. Population structure of *Escherichia coli* causing bacteraemia in the UK and Ireland between 2001 and 2010. J Antimicrob Chemother. 2016;71: 2139–2142.

50. Bortolaia V, Larsen J, Damborg P, Guardabassi L. Potential pathogenicity and host range of extended-spectrum beta-lactamase-producing *Escherichia coli* isolates from healthy poultry. Appl Environ Microbiol. 2011;77: 5830–5833.

51. Letunic I, Bork P. Interactive tree of life (iTOL) v3: an online tool for the display and annotation of phylogenetic and other trees. Nucleic Acids Res. 2016;44: W242–5.

52. Selander RK, Caugant DA, Whittam TS. Genetic structure and variation in natural populations of Escherichia coli. 1987.

53. Herzer PJ, Inouye S, Inouye M, Whittam TS. Phylogenetic distribution of branched RNA-linked multicopy single-stranded DNA among natural isolates of *Escherichia coli*. J Bacteriol. 1990;172: 6175–6181.

54. Clermont O, Olier M, Hoede C, Diancourt L, Brisse S, Keroudean M, et al. Animal and human pathogenic *Escherichia coli* strains share common genetic backgrounds. Infect Genet Evol. 2011;11: 654–662.

55. Clermont O, Dixit OVA, Vangchhia B, Condamine B, Dion S, Bridier-Nahmias A, et al. Characterization and rapid identification of phylogroup G in *Escherichia coli*, a lineage with high virulence and antibiotic resistance potential. Environ Microbiol. 2019;21: 3107–3117.

56. Waters NR, Abram F, Brennan F, Holmes A, Pritchard L. Easily phylotyping *E. coli* via the EzClermont web app and command-line tool. Access Microbiology. 2020;6: acmi000143

57. Baker KS, Burnett E, McGregor H, Deheer-Graham A, Boinett C, Langridge GC, et al. The Murray collection of pre-antibiotic era Enterobacteriacae: a unique research resource. Genome Med. 2015;7: 97.

58. Magiorakos A-P, Srinivasan A, Carey RB, Carmeli Y, Falagas ME, Giske CG, et al. Multidrug-resistant, extensively drug-resistant and pandrug-resistant bacteria: an international expert proposal for interim standard definitions for acquired resistance. Clin Microbiol Infect. 2012;18: 268–281.

59. Touchon M, Perrin A, de Sousa JAM, Vangchhia B, Burn S, O’Brien CL, et al. Phylogenetic background and habitat drive the genetic diversification of *Escherichia coli*. PLoS Genet. 2020;16: e1008866.

60. Hazen TH, Donnenberg MS, Panchalingam S, Antonio M, Hossain A, Mandomando I, et al. Genomic diversity of EPEC associated with clinical presentations of differing severity. Nat Microbiol. 2016;1: 15014.

61. Burger R. EHEC O104:H4 in Germany 2011: Large outbreak of bloody diarrhea and haemolytic uraemic syndrome by shiga toxin-producing *E. coli* via contaminated food. National Academies Press (US); 2012.

62. Camacho C, Coulouris G, Avagyan V, Ma N, Papadopoulos J, Bealer K, et al. BLAST+: architecture and applications. BMC Bioinformatics. 2009;10: 421.

63. Buchfink B, Xie C, Huson DH. Fast and sensitive protein alignment using DIAMOND. Nat Methods. 2015;12: 59–60.

64. von Mentzer A, Connor TR, Wieler LH, Semmler T, Iguchi A, Thomson NR, et al. Identification of enterotoxigenic *Escherichia coli* (ETEC) clades with long-term global distribution. Nat Genet. 2014;46: 1321–1326.

65. Kallonen T, Brodrick HJ, Harris SR, Corander J, Brown NM, Martin V, et al. Systematic longitudinal survey of invasive *Escherichia coli* in England demonstrates a stable population structure only transiently disturbed by the emergence of ST131. Genome Res. 2017. doi: 10.1101/gr.216606.116

66. Salipante SJ, Roach DJ, Kitzman JO, Snyder MW, Stackhouse B, Butler-Wu SM, et al. Large-scale genomic sequencing of extraintestinal pathogenic *Escherichia coli* strains. Genome Res. 2015;25: 119–128.

67. Abram KZ, Udaondo Z, Bleker C, Wanchai V. What can we learn from over 100,000 *Escherichia coli* genomes? bioRxiv. 2020. Available: https://www.biorxiv.org/content/10.1101/708131v2.abstract

68. Ingle DJ, Tauschek M, Edwards DJ, Hocking DM, Pickard DJ, Azzopardi KI, et al. Evolution of atypical enteropathogenic *E. coli* by repeated acquisition of LEE pathogenicity island variants. Nat Microbiol. 2016;1: 15010.

69. Hazen TH, Sahl JW, Fraser CM, Donnenberg MS, Scheutz F, Rasko DA. Refining the pathovar paradigm via phylogenomics of the attaching and effacing *Escherichia coli*. Proc Natl Acad Sci U S A. 2013;110: 12810–12815.

70. Goh KGK, Phan M-D, Forde BM, Chong TM, Yin W-F, Chan K-G, et al. Genome-Wide Discovery of Genes Required for Capsule Production by Uropathogenic *Escherichia coli*. MBio. 2017;8. doi: 10.1128/mBio.01558-17

71. Chen SL, Wu M, Henderson JP, Hooton TM, Hibbing ME, Hultgren SJ, et al. Genomic diversity and fitness of *E. coli* strains recovered from the intestinal and urinary tracts of women with recurrent urinary tract infection. Sci Transl Med. 2013;5: 184ra60.

