## Supplementary for "A comprehensive and high-quality collection of *E. coli* genomes and their genes": S1.pdf

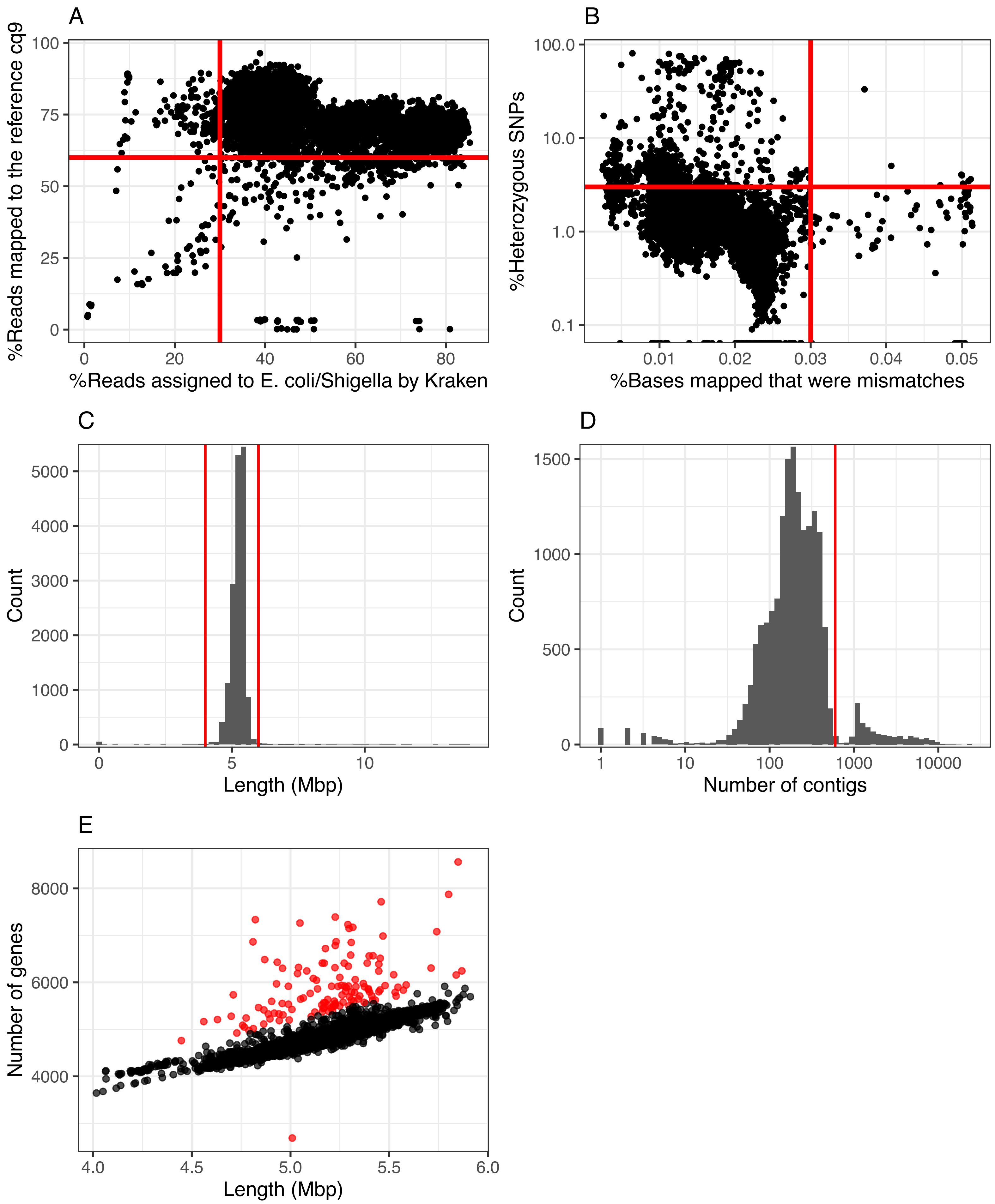


**S1: Quality control measures used to filter genomes.A** Percentage of reads which were assigned to *E. coli/Shigella* using Kraken relative to the number of reads mapped to an *E. coli* reference cq9. Red lines indicate cut-offs applied, top right corner are all remaining genomes. **B** Percentage of bases mapped which were mismatches relative to the percentage of heterozygous SNPs for each genome. Red lines indicate cut-offs applied, bottom left corner are all remaining genomes. **C** Distribution of genome lengths in the collection. Red lines: genomes shorter than 4 Mb or longer than 6 Mb were removed. **D** Distribution of number of contigs per genome in the collection. Red line: genomes with more than 600 contigs were removed. **E** Correlation between genome length and number of predicted CDSs using Prokka. Red: Genomes which deviated from the expected number of genes were removed.
