## Supplementary for "A comprehensive and high-quality collection of *E. coli* genomes and their genes": S2.pdf

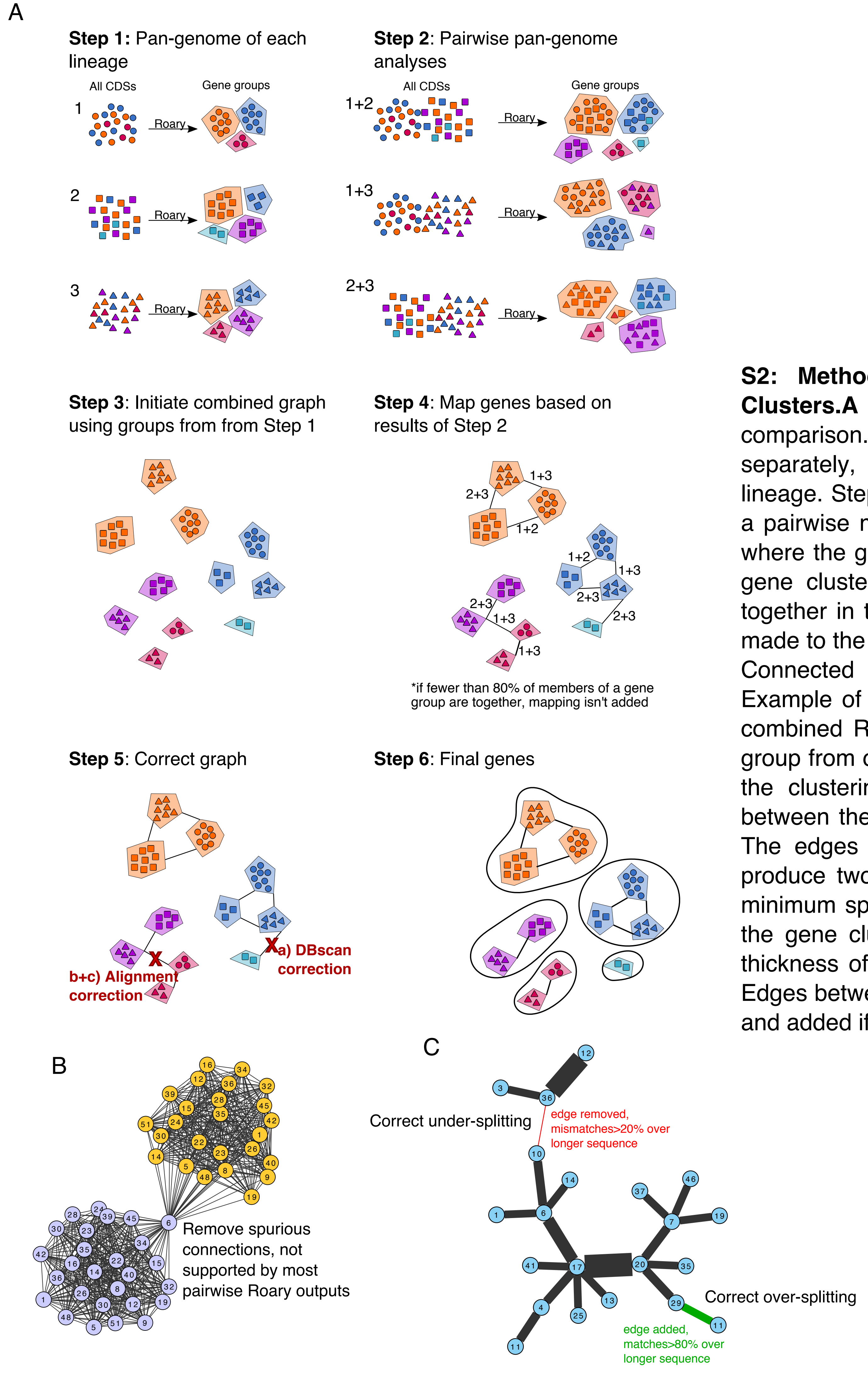


**S2: Method for combining the pan-genome analysis of all PopPUNK Clusters.A** Procedure for a pan-genome analysis using a pairwise Roary comparison. Step 1: a pan-genome analysis was applied on each lineage separately, generating gene clusters from all the CDSs of all genomes in that lineage. Step 2: A pan-genome analysis using Roary was applied on all lineages in a pairwise manner, generating new gene clusters. Step 3: A graph is constructed where the gene clusters from Step 1 are the nodes. Step 4: An edge between two gene clusters was added if the members of both gene clusters were grouped together in the pairwise pan-genome analysis in Step 2. Step 5: Corrections were made to the graph using density based clustering and sequence alignments. Step 6: Connected components were extracted as the final gene cluster definitions. **B** Example of density based clustering correction. The graph is a real example of a combined Roary graph as presented at the end of Step 4. Each node is a gene group from one lineage. The nodes are numbered by their Lineage and coloured by the clustering result of density based clustering. In this case, the connection between the two groups is only supported by a spurious connection of Lineage 6. The edges between Lineage 6 and the rest of yellow lineage are removed to produce two groups. **C** Example of alignment based corrections. The graph is a minimum spanning tree of the alignment between the representative sequences of the gene clusters from each lineage. Each node is a gene in one lineage. The thickness of the edge between two genes in the percent matches between them. Edges between genes are removed if they differ by more than 20% (under-splitting), and added if they match by more than 80% (over-splitting).
