## Supplementary for "A comprehensive and high-quality collection of *E. coli* genomes and their genes": S3.pdf

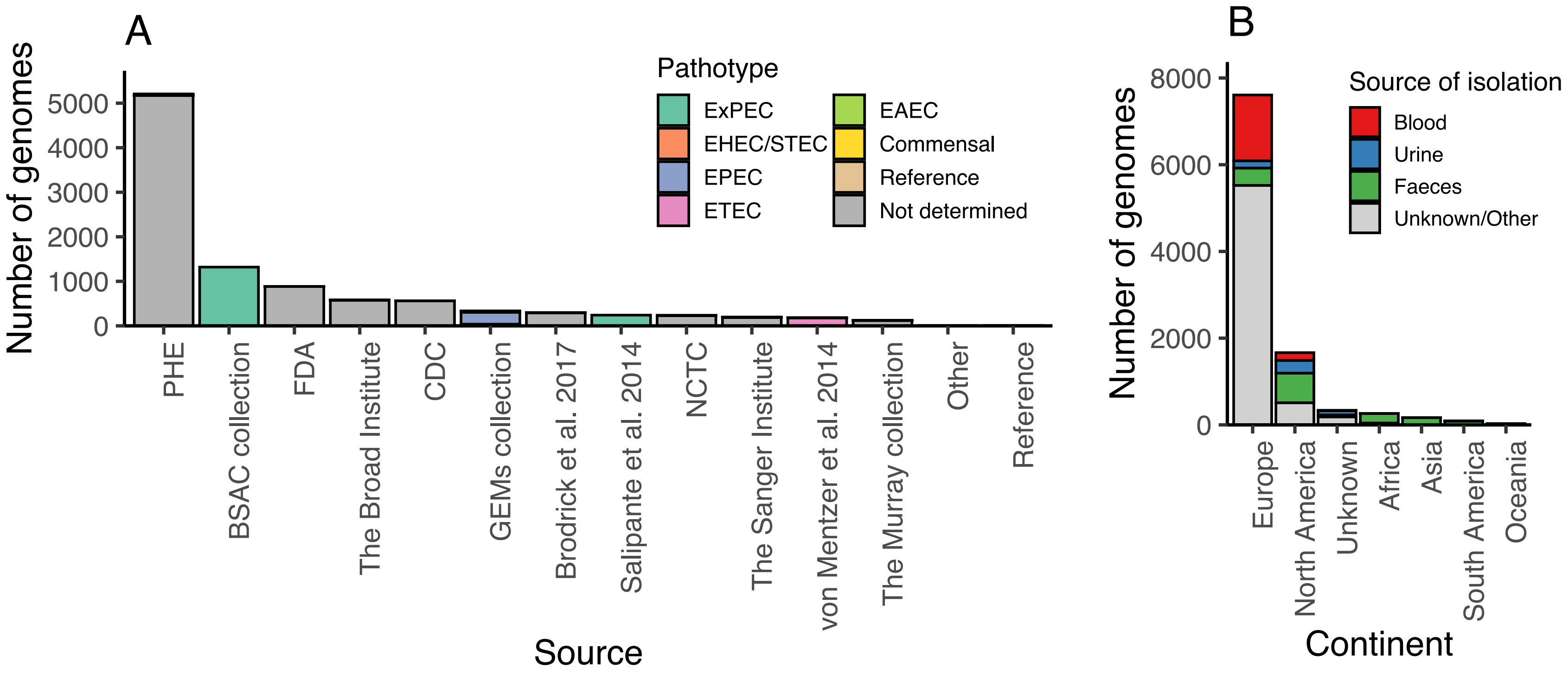


**S3: Source of *E. coli* genomes. A** Source of the *E. coli* genomes in the collection, coloured by the pathotype associated with the specific studies. **B** Continents from which the *E. coli* genomes were collected, coloured by source of isolation.
