## Supplementary for "A comprehensive and high-quality collection of *E. coli* genomes and their genes": S4.pdf

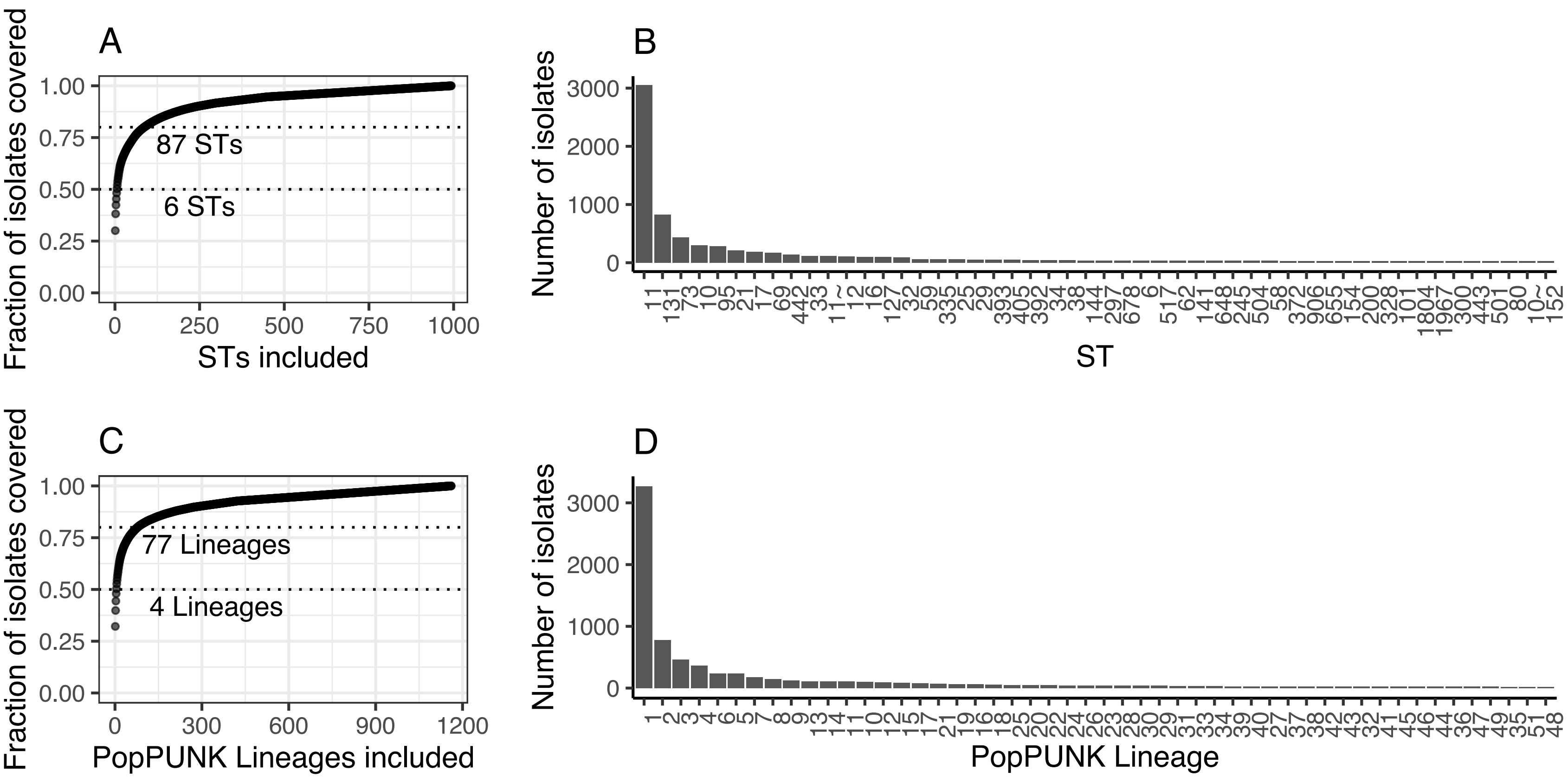


**S4: Distribution of STs and PopPUNK Lineages in the collection. A,C** Coverage of genome collection by increasing the number of STs (A) or PopPUNK Lineages (C) included in the study. Dotted lines: Number of STs (A) or PopPUNK Lineages (C) which accounted for 0.5 and 0.8 of all isolates in the genome collection. **B,D** Number of genomes in the fifty largest STs (B) and PopPUNK Lineages (D).
