## Supplementary for "A comprehensive and high-quality collection of *E. coli* genomes and their genes": S5.pdf

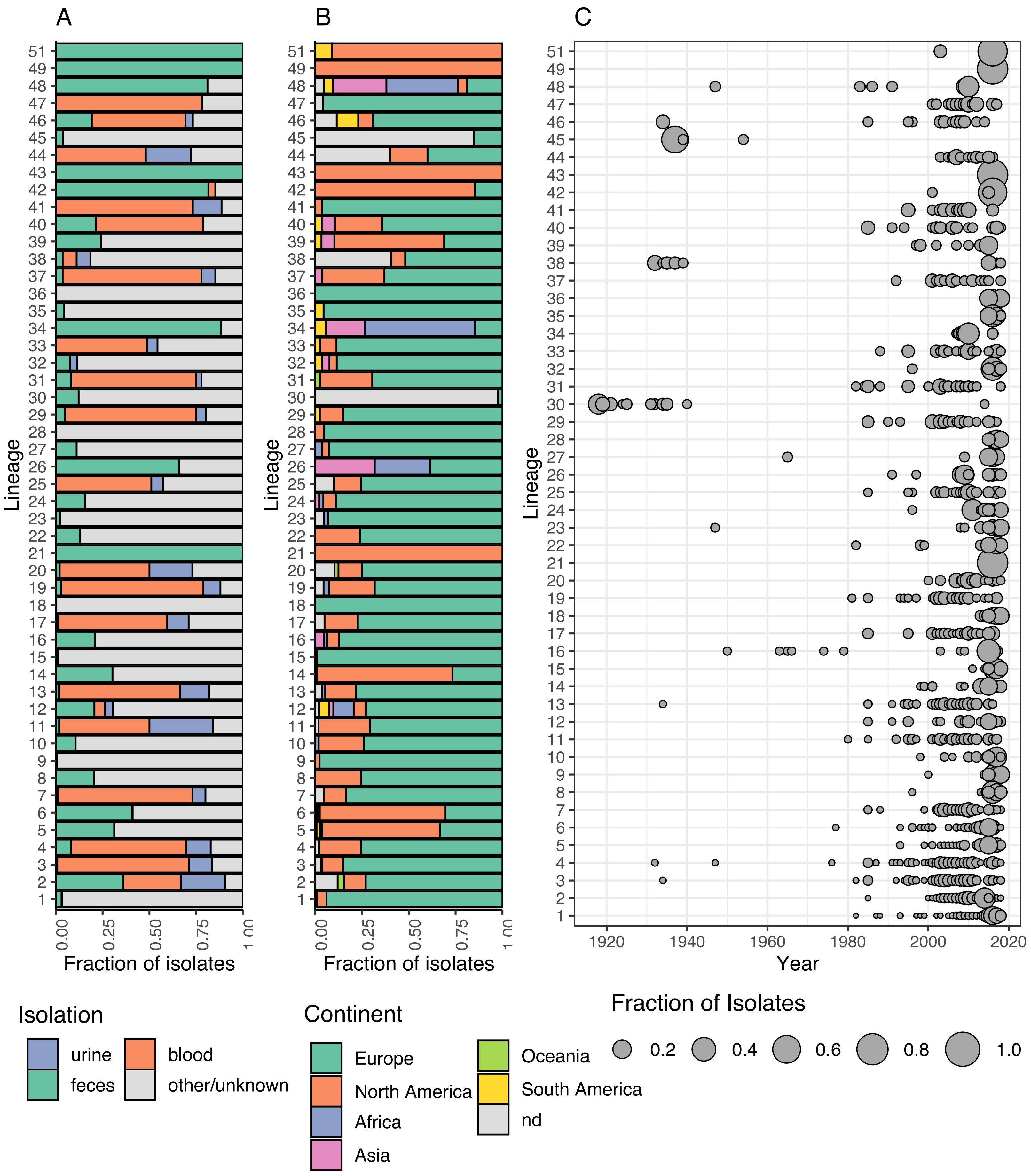


**S5: Metadata associated with the lineages.A,B** Source of isolation (A) and continent (B) of isolates from the fifty lineages. **C** Fraction of genomes from each of the lineages collected from each year (where metadata was available).
