## Supplementary for "A comprehensive and high-quality collection of *E. coli* genomes and their genes": S6.pdf

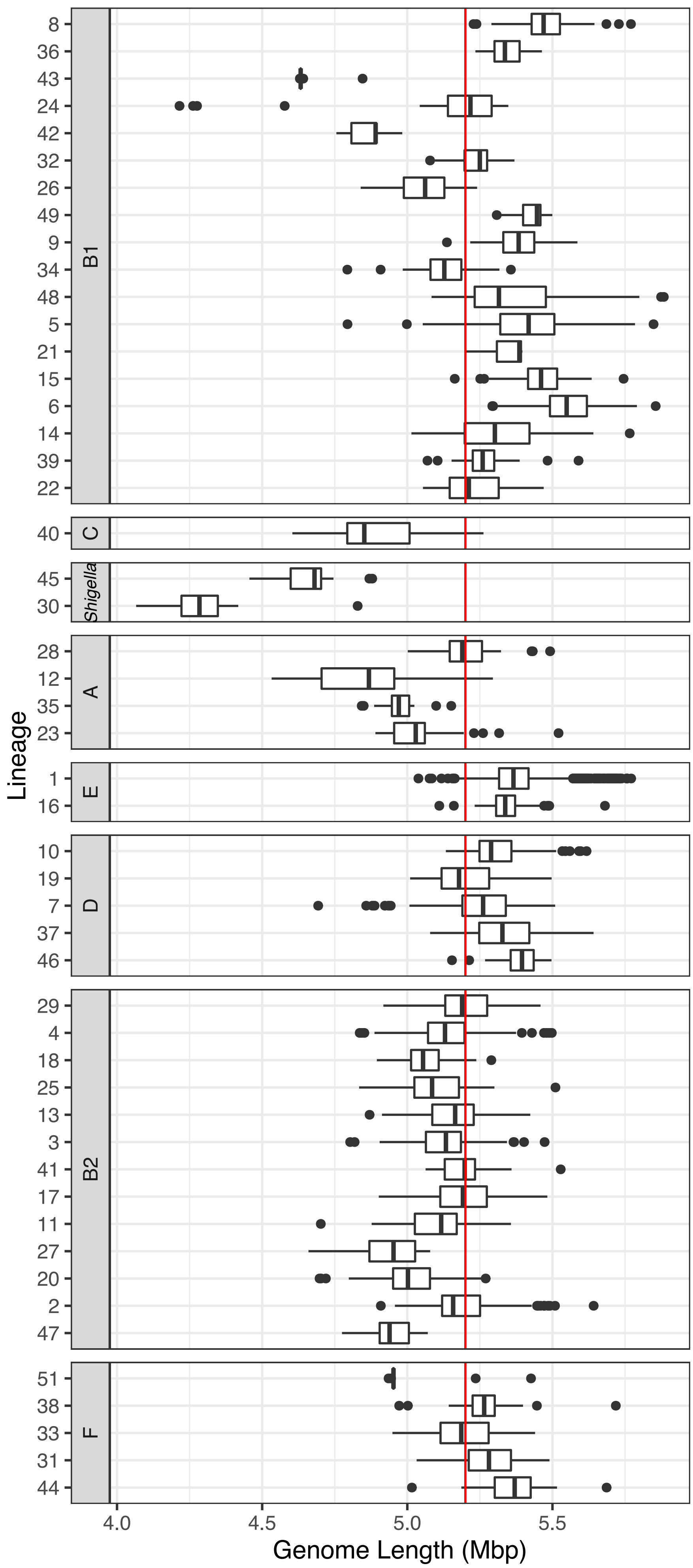


**S6: Genome lengths across the lineages.** Genome length, measured as total contig length, per isolate across the lineages, divided by their phylogroup. Red line: weighted-mean genome length across the entire collection.
