## Supplementary for "A comprehensive and high-quality collection of *E. coli* genomes and their genes": S7.pdf

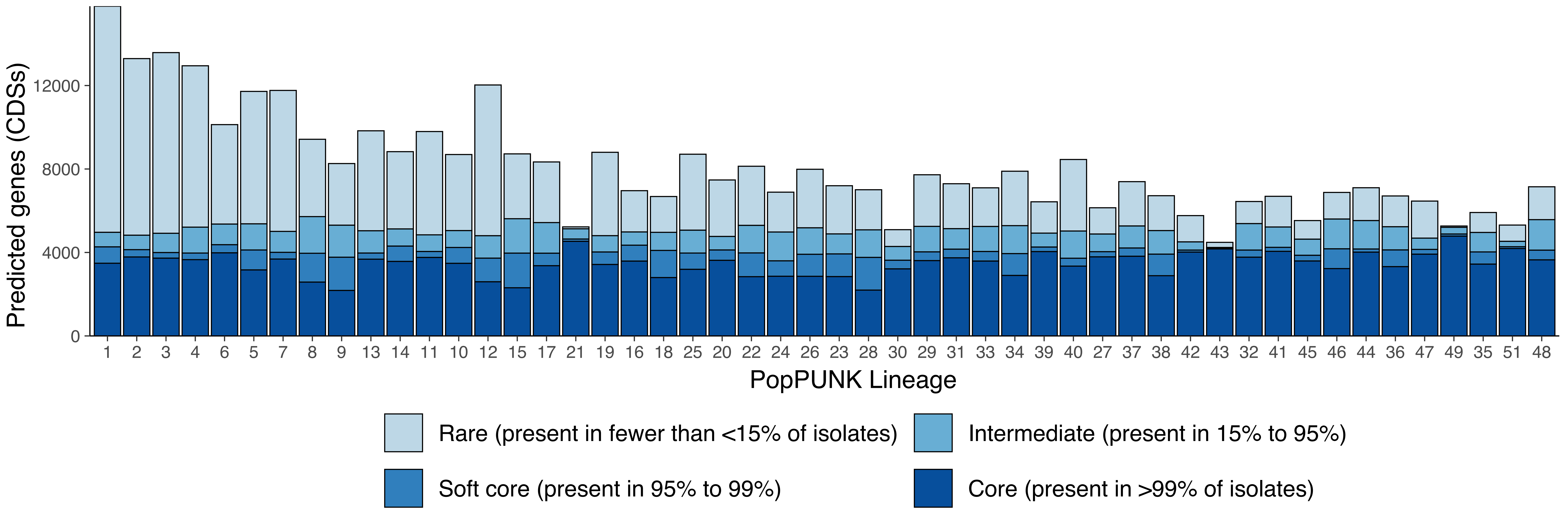


**S7: Pan-genome size across the lineages.** Number of predicted CDSs in each lineage in a Roary analysis per lineage, coloured by division into core, soft-core, intermediate and rare genes. Lineages 21, 43 and 49 were removed from downstream analysis due to low diversity in gene content.
